# Misexpression of inactive genes in whole blood is associated with nearby rare structural variants

**DOI:** 10.1101/2023.11.17.567537

**Authors:** Thomas Vanderstichele, Katie L Burnham, Niek de Klein, Manuel Tardaguila, Brittany Howell, Klaudia Walter, Kousik Kundu, Jonas Koeppel, Wanseon Lee, Alex Tokolyi, Elodie Persyn, Artika P Nath, Jonathan Marten, Slavé Petrovski, David J Roberts, Emanuele Di Angelantonio, John Danesh, Alix Berton, Adam Platt, Adam S Butterworth, Nicole Soranzo, Leopold Parts, Michael Inouye, Dirk S Paul, Emma E Davenport

## Abstract

Gene misexpression is the aberrant transcription of a gene in a context where it is usually inactive. Despite its known pathological consequences in specific rare diseases, we have a limited understanding of its wider prevalence and mechanisms in humans. To address this, we analyzed gene misexpression in 4,568 whole blood bulk RNA sequencing samples from INTERVAL study blood donors. We found that while individual misexpression events occur rarely, in aggregate they were found in almost all samples and over half of inactive genes. Using 2,821 paired whole genome and RNA sequencing samples, we identified that misexpression events are enriched in *cis* for rare structural variants. We established putative mechanisms through which a subset of SVs lead to gene misexpression, including transcriptional readthrough, transcript fusions and gene inversion. Overall, we develop misexpression as a novel type of transcriptomic outlier analysis and extend our understanding of the variety of mechanisms by which genetic variants can influence gene expression.

## Introduction

Temporal and spatial regulation of gene expression is essential for the functioning of multicellular eukaryotes. Gene regulation involves the context-specific activation and maintenance of transcription, as well as gene silencing to avoid aberrant transcription interfering with normal cellular function. The aberrant transcription of a gene in a context where it is usually inactive is termed gene misexpression (also referred to as ectopic expression) (**Fig.1A**)^1^. Gene misexpression can occur either via the transcription of a single inactive gene or production of a novel transcript derived in part from an inactive gene. We refer to these different types of events as non-chimeric and chimeric misexpression, respectively.

**Figure 1.**
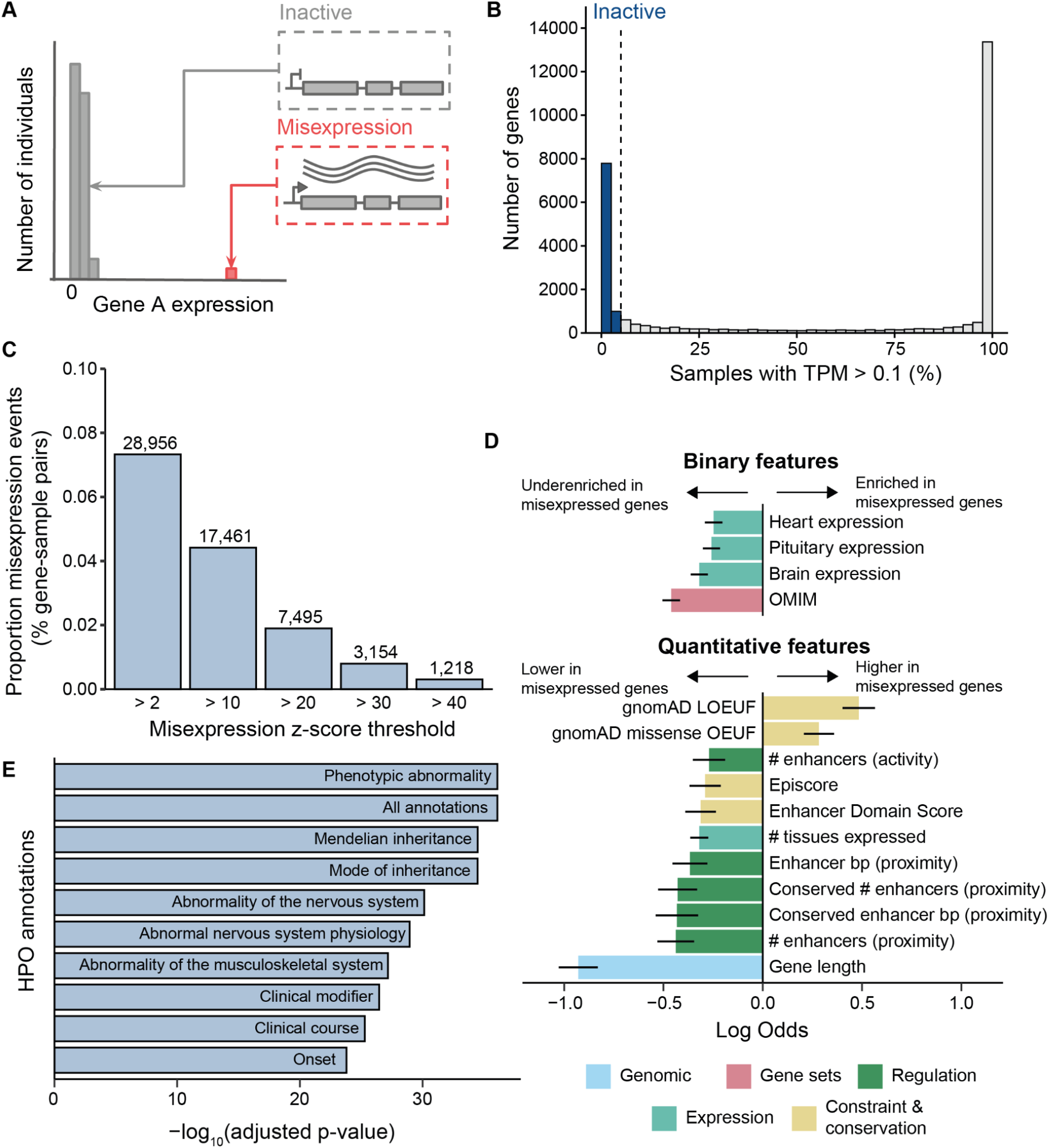
Identification of misexpression events and characterisation of misexpressed genes. **A.)** Gene misexpression is the aberrant transcription of a gene in a context where it is usually inactive. In this schematic, the majority of individuals have negligible or no expression of gene A (inactive, gray) with only a handful of individuals showing high expression (misexpression, red). **B.)** Distribution of gene activity across 29,614 genes within the INTERVAL whole blood RNA-seq dataset. For each gene, activity is quantified as the percentage of samples where the gene has a TPM > 0.1 (x-axis). Inactive genes are defined as having a TPM > 0.1 in less than 5% of samples (vertical dashed line). **C.)** Proportion of 39,513,200 gene-sample pairs (8,650 inactive genes across 4,568 samples) that are misexpressed (y-axis) across different misexpression z-score thresholds (x-axis). Text labels indicate the total number of misexpression events at each misexpression z-score threshold. **D.)** Enrichment of gene-level features within 4,437 genes that are misexpressed (z-score > 2 and TPM > 0.5) versus 4,213 non-misexpressed genes. The 15 features with the highest absolute log odds passing Bonferroni correction are shown. Lines indicate 95% confidence intervals for the fitted parameters using the standard normal distribution. **E.)** Top 10 Human Phenotype Ontology (HPO) terms, by -log_10_(adjusted p-value) on the x-axis, underrepresented within 4,437 misexpressed genes using all 8,650 inactive genes as the custom background. OMIM; Online Mendelian Inheritance in Man, LOEUF; loss-of-function observed/expected upper bound fraction, OEUF; observed/expected upper bound fraction.

Gene misexpression can have profound phenotypic consequences, as evidenced by the development of ectopic eyes across different tissues in *Drosophila melanogaster* upon targeted misexpression of the *eyeless* gene^2^. In humans, gene misexpression has been implicated in cancers^3,4^ and several rare diseases, for example, congenital limb malformations^5^, congenital hyperinsulinism^6^ and monogenic severe childhood obesity^7^. These studies have identified gain-of-function genetic variants that lead to both chimeric and non-chimeric gene misexpression. For example, chimeric misexpression can be caused by transcript fusions^7^ and non-chimeric misexpression via rearrangements in 3D chromatin architecture^8^ or loss of silencer function^6^. However, these studies have predominantly focused on a limited number of disease-related loci.

Recent large-scale RNA sequencing (RNA-seq) studies analyzing transcriptional outliers in humans have demonstrated that outliers are enriched for rare single nucleotide variants (SNVs), indels and structural variants (SVs) in *cis^9–12^* and that these outlier-associated genetic variants can contribute to complex disease risk^11,13^. However, these studies focused on outliers in highly expressed genes within the tissue(s) under study, overlooking misexpression of inactive genes. Consequently, the prevalence of gene misexpression in humans, the genes whose misexpression can be tolerated and their associated properties are unknown. Furthermore, the types of genetic variants associated with misexpression and their mechanisms remain underexplored.

To address these gaps in our understanding, we conducted a genome-wide analysis of gene misexpression using bulk RNA-seq data from 4,568 blood donors from the INTERVAL study^14,15^. We assessed the prevalence of gene misexpression across genes and samples, and the characteristics of genes that tolerate misexpression. Additionally, we established the types of genetic variants associated with gene misexpression as well as their putative mechanisms of action using 2,821 paired whole genome sequencing (WGS) and RNA-seq samples.

## Results

### Identification of misexpression events in whole blood

To identify misexpression events, we first defined a set of inactive genes with negligible or no detectable expression across the majority of the 4,568 whole blood RNA-seq samples from the INTERVAL study (**Methods**). We restricted our analysis to 29,614 autosomal protein-coding and long non-coding RNA genes with evidence of being expressed in at least one tissue from the Genotype-Tissue Expression (GTEx) project. From these, we identified 8,779 inactive genes that were expressed (transcripts per million (TPM) > 0.1) in less than 5% of samples (**Fig. 1B**). We confirmed that these genes were likely inactive using other whole blood RNA-seq datasets, such as GTEx, and predicted chromatin states from peripheral blood mononuclear cells (PBMCs) (**Supplementary Fig. 1**). To account for non-genetic drivers of misexpression, such as sequencing depth or variation in cell proportions, we removed 129 (1.5%) genes that were significantly correlated (|Spearman’s rho| > 0.2, FDR-adjusted p < 0.05) with any of 225 technical and cellular covariates. We transformed expression values into z-scores and identified 28,956 misexpression events (z-score > 2 and TPM > 0.5). Across all inactive gene-sample pairs, the proportion of misexpression events was low (0.07%, 28,956/39,513,200) with the number of events decreasing substantially at higher z-score thresholds (**Fig. 1C, Supplementary Fig. 2**). While individual misexpression events occurred rarely, in aggregate they were found in 51% of inactive genes (4,437/8,650) and in 96% of samples (4,386/4,568) with a median of 4 events per sample (**Supplementary Fig. 2**).

### Misexpressed genes are shorter, depleted of developmental genes and less tightly regulated

Next, we investigated the properties that differ between genes with and without misexpression events. We tested for enrichment of 81 gene-level features in genes that were misexpressed at least once (’’misexpressed genes’’) versus genes with no observed misexpression events (’’non-misexpressed genes’’) across different misexpression z-score thresholds (**Fig. 1D, Supplementary Fig. 3, Methods**). Overall, misexpressed genes were shorter, less constrained according to both mutational (gnomAD LOEUF and missense OEUF) and non-mutational (Enhancer Domain Score and Episcore) metrics and were less likely to be implicated in developmental diseases^16–19^. These genes also had fewer predicted enhancer interactions based on both proximity- and activity-linking approaches from Wang &

Goldstein^17^, suggesting that they are under weaker regulatory control. Misexpressed genes were less likely to be expressed in brain, pituitary and heart tissues, and generally were expressed across fewer GTEx tissues (n = 52). Additionally, they were underrepresented for Human Phenotype Ontology (HPO) terms relating to phenotypic abnormalities of the nervous and musculoskeletal system (**Fig. 1E**, **Supplementary Fig. 4, Methods**). This is of interest as congenital limb malformations are known to be caused by gene misexpression^8^. Taken together, these results suggest that natural selection has acted to prevent the misexpression of genes important in developmental processes. Importantly, these results also demonstrate that our method for identifying gene misexpression is valid, as we would expect misexpression of inactive developmental genes to be deleterious, and therefore, underenriched in a generally healthy population cohort.

### Rare structural variants are associated with gene misexpression

To assess the influence of genetic variation on gene misexpression in *cis* we conducted genetic variant enrichment analyses. Our analysis focused on 2,821 participants with both WGS and RNA-seq data in the INTERVAL study (**Methods**). In total, we conducted 700 enrichment tests, determining significance using a Bonferroni-adjusted p-value threshold (p < 0.05). Firstly, we tested whether rare (MAF < 1%), low-frequency (1% ≤ MAF < 5%) or common (5% ≤ MAF < 50%) SNVs, indels (≤ 50 bp) or SVs (> 50 bp) were enriched within the gene body and flanking sequence of genes involved in misexpression events (±10 kb for SNVs and indels, ±200 kb for SVs, **Methods**). Across all tested z-score thresholds, we observed a significant enrichment of rare SVs around gene misexpression events, whereas no significant enrichment was observed for low frequency or common SVs at any z-score threshold (**Fig. 2A**). The enrichment for rare SVs increased dramatically at increasing z-score thresholds, with 4.7% (38/803) of extreme misexpression events (z-score > 40) having a nearby rare SV compared to 1.3% (229/17,380) of less extreme events (z-score > 2, **Fig. 2A**, **Supplementary Fig. 5**). Notably, we did not find a significant enrichment for SNVs or indels at any MAF or z-score threshold (maximum enrichment SNVs = 1.04 and indels = 1.15) and even observed a significant weak underenrichment in some cases.

**Figure 2.**
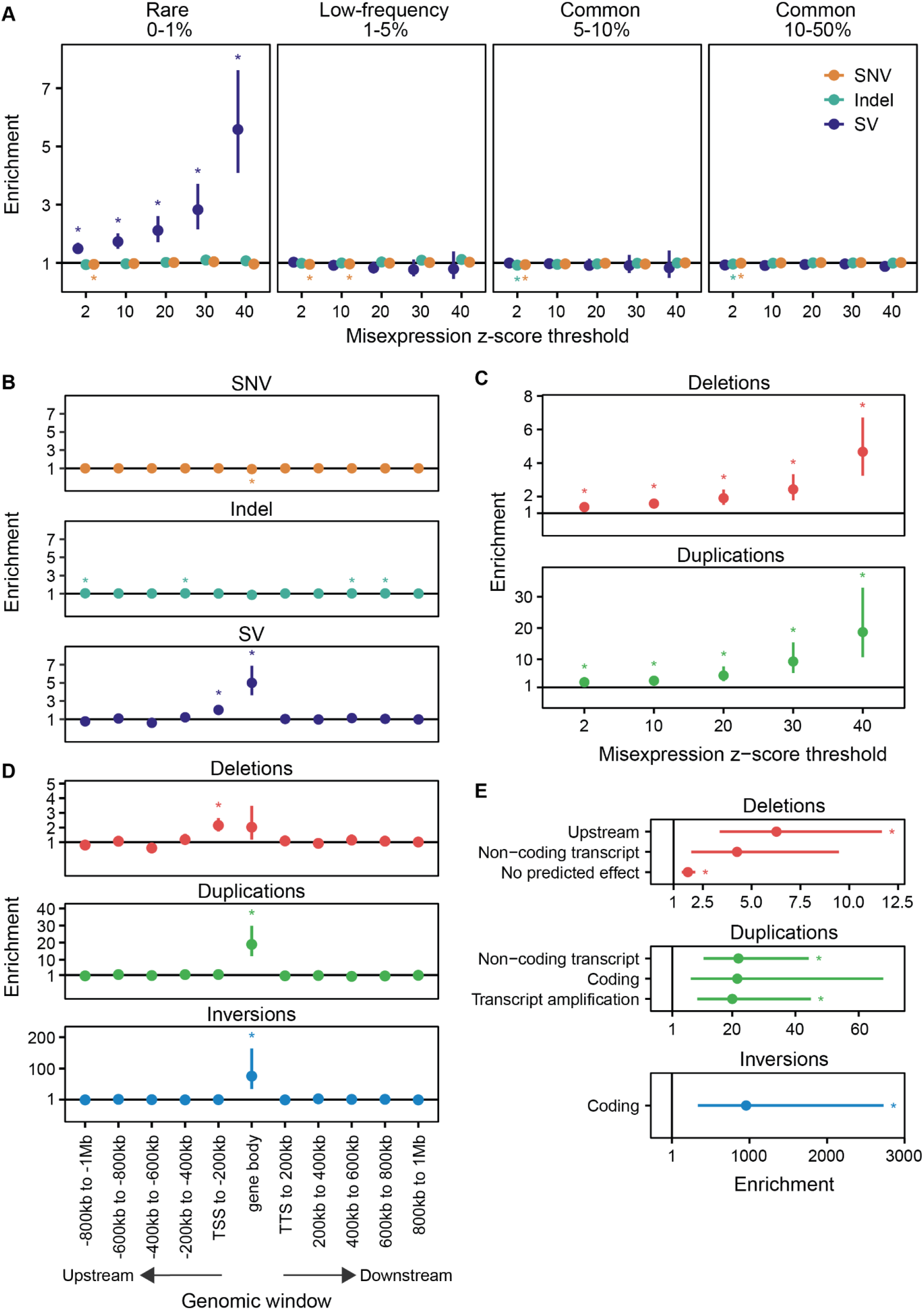
Enrichment of genetic variants near to gene misexpression events. Across all figures, enrichments were calculated as the relative risk of having a nearby variant type or consequence given the misexpression status. Bars represent 95% Wald confidence intervals of the relative risk estimates. The line at enrichment = 1 indicates no enrichment; asterisks positioned either side of the line indicate significant enrichment or underenrichment after Bonferroni correction. **A.)** Enrichment of SNVs, indels and SVs within the gene body and flanking sequence of genes involved in misexpression events across different misexpression z-score thresholds and MAF cutoffs. A flanking sequence of ±200 kb around each gene was used for SVs and ±10 kb for SNVs and indels. **B.)** Enrichment of rare (MAF < 1%) SNVs, indels and SVs within 200 kb genomic windows and the body of genes involved in misexpression events. The misexpression threshold shown is a z-score > 10 and TPM > 0.5. **C.)** Enrichment of rare (MAF < 1%) deletions and duplications in a ±200 kb window around genes involved in misexpression events across different misexpression z-score thresholds. **D.)** Enrichment of rare (MAF < 1%) deletions, duplications, and inversions within 200 kb genomic windows and the body of genes involved in misexpression events. The misexpression threshold shown is a z-score > 10 and TPM > 0.5. **E.)** Enrichment of rare (MAF < 1%) SVs, stratified by their class and predicted VEP consequence in a ±200 kb window around genes involved in misexpression events. The misexpression threshold shown is a z-score > 10 and TPM > 0.5. Only SV consequences with at least one Bonferroni significant enrichment at any z-score threshold are shown. SNV; single nucleotide variant, SV; structural variant, TTS; transcription termination site, TSS; transcription start site.

We examined whether the observed rare SV enrichment could be due to a small number of SVs leading to the misexpression of many genes. However, out of the 312 SVs within 200kb of a misexpression event, 95% (297) were linked to only one gene and the remainder to a maximum of two genes. We also assessed whether the observed rare SV enrichment could be due to a small number of participants with a high number of SVs and misexpression events. Similarly, of the 206 participants containing misexpression events with a nearby SV, 89% (183) had only one misexpression event with an SV in *cis*.

To investigate the influence of genetic variation on gene misexpression over longer distances, we tested rare variant enrichment at increasing distances from genes. For each gene, we assigned each rare SNV, indel and SV to a unique genomic window up to 1 Mb upstream or downstream and tested variant enrichment for each window independently (**Methods**). Across all z-score thresholds, enrichment was highest for rare SVs within the gene body and decreased at greater distances from the misexpressed gene (**Fig. 2B**, **Supplementary Fig. 6**), remaining significant up to 200 kb upstream. Interestingly, rare SV enrichment was not symmetrical around misexpressed genes with greater enrichment upstream of transcription start sites (TSS) compared to downstream of the transcription termination site (TTS). Similarly to the gene-level analysis, we found that rare SNVs were not significantly enriched across any window or expression threshold, and again observed significant weak underenrichment in some genomic windows (**Fig. 2B**, **Supplementary Fig. 6**). While indels did show a significant enrichment in some windows, the level of enrichment was much lower than SVs (maximum significant enrichment = 1.11) and was not consistently observed across all z-score thresholds (**Supplementary Fig. 6**).

### Rare deletions, duplications and inversions are associated with gene misexpression

We hypothesized that misexpression events are associated with a specific type of structural variation and therefore conducted enrichment tests for the four different SV classes available: deletions (DEL), duplications (DUP), inversions (INV) and mobile element insertions (MEI) (**Fig. 2C**, **Supplementary Fig. 7**, **Methods**). Across all z-score thresholds, rare deletions and duplications were significantly enriched within a 200 kb window around misexpressed genes with duplications consistently showing the highest enrichment. However, at this sample size and genomic window size, rare inversions and mobile element insertions were not significantly enriched.

Next, we tested whether different SV classes showed distinct patterns of enrichment at increasing distances from misexpressed genes (**Methods**). Rare duplications and inversions were significantly enriched only within the gene body of the misexpressed gene (**Fig. 2D**). For duplications, all tested z-score thresholds were significant, while for inversions this enrichment was significant only up to a z-score threshold of 10, likely due to the low number of inversion calls (**Supplementary Fig. 7**). Rare deletions were significantly enriched in the window 200 kb upstream from the transcription start site (TSS) across all z-score thresholds (**Fig. 2D**, **Supplementary Fig. 7**). Rare deletions were also enriched within the gene body of the misexpressed gene but this was only significant at higher z-score thresholds (**Supplementary Fig. 7**). Rare mobile element insertions were not significantly enriched within any tested window at any z-score threshold. No significant enrichment was observed at greater distances for any SV class.

We annotated each rare SV by its predicted consequence on the inactive genes in the tested window using the Ensembl Variant Effect Predictor (VEP)^20^. For each SV class, we then tested for enrichment of predicted consequences ±200 kb around misexpressed genes relative to controls (**Fig. 2E**, **Supplementary Fig. 8**, **Methods**). We found that inversions affecting coding regions had the highest enrichment of any variant consequence; however, this was only significant up to a z-score threshold of 20, again likely due to the low number of inversion calls. Deletions upstream and with no predicted effect on the tested gene were significantly enriched as were deletions affecting non-coding transcripts at higher z-score thresholds. Additionally, duplications leading to transcript amplification and affecting coding regions as well as non-coding transcripts were significantly enriched. These results support the enrichment observed for SVs classes within specific genomic windows.

### Properties and regulatory features of misexpression-associated structural variants

To understand the general properties of misexpression-associated rare SVs, we compared a set of 105 misexpression-associated and 20,150 control SVs (**Methods**). Of these 105 SVs, 87 were deletions, 16 were duplications and 2 were inversions (**Fig. 3A**). All the duplications were confirmed to be tandem duplications (**Methods**). Notably, for 60% and 28% of deletions and duplications VEP did not predict an effect on the misexpressed gene (**Fig. 3B**). While the majority (72%) of duplications overlapped the misexpressed gene either entirely or partially, this was not the case for deletions (8% overlapping) (**Fig. 3C**). Therefore, we analyzed the properties of deletions and duplications separately, excluding inversions due to their low numbers.

**Figure 3.**
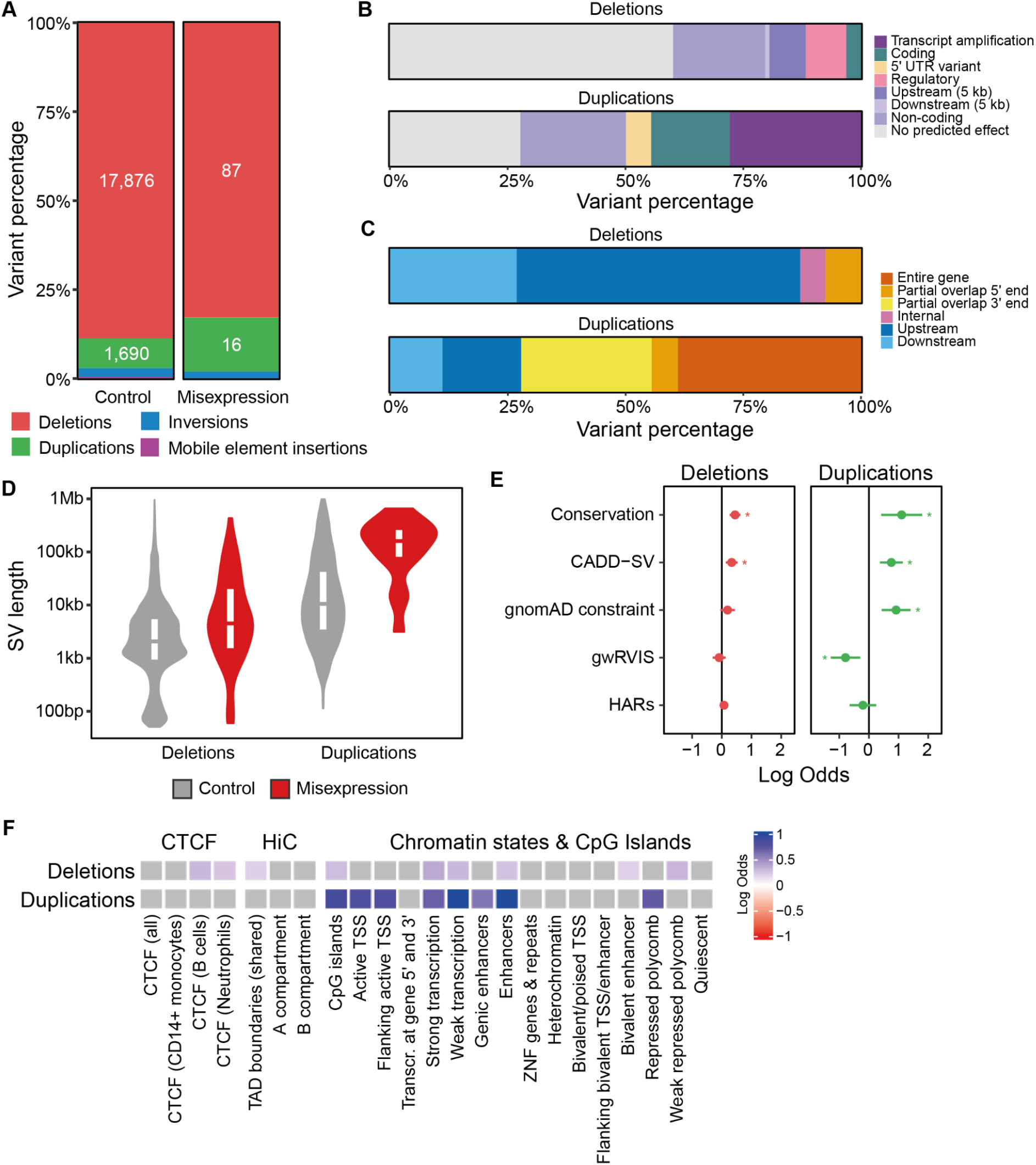
Properties and regulatory features associated with misexpression-associated rare SVs. **A.)** Proportion of misexpression-associated and control deletions, duplications, inversions and mobile element insertions. Proportion of misexpression-associated deletions and duplications by their **B.)** predicted VEP consequence on the misexpressed gene and **C.)** position relative to the misexpressed gene. **D.)** SV length distributions of misexpression-associated and control duplications and deletions. **E.)** Enrichment (x-axis) of misexpression-associated deletions (left panel, red) and duplications (right panel, green) compared to controls for genomic scores (y-axis) including evolutionary conservation (PhyloP), predicted deleteriousness (CADD-SV), constraint (gnomAD z-score constraint and gwRVIS), and HARs. Enrichments were calculated as the log odds ratio with lines indicating 95% confidence intervals for the fitted parameters using the standard normal distribution. Asterisks indicate significant enrichment after Bonferroni correction. **F.)** Enrichment of misexpression-associated deletions and duplications compared to controls for regulatory features including CTCF candidate cis-regulatory elements (cCREs) from ENCODE, TAD boundaries shared across multiple cell-lines, A and B compartments, chromatin states from the Roadmap Epigenomics Project and CpG islands from the UCSC genome browser. Enrichments were calculated as the log odds ratio and tiles shaded in gray do not pass Bonferroni correction. CADD-SV; combined annotation dependent depletion for SVs, gwRVIS; genome-wide residual variation intolerance score, TADs; topologically-associated domains, HARs; human accelerated regions, TSS; transcription start site, CTCF; CCCTC-binding factor, ZNFs; zinc-finger proteins.

First, we found that misexpression-associated deletions and duplications were on average 2.5- and 3.4-times longer, respectively, than control variants (p = 6.03×10^-7^ and p = 3.9×10^-6^, one-sided Mann-Whitney U test, **Fig. 3D**). Since MAF and SV length are inversely correlated, we also compared the lengths of singletons and found that misexpression-associated deletions and duplications remained on average 2.8- and 2.0-times longer, respectively (p = 2.3×10^-4^ and p = 2.2×10^-3^, one-sided Mann-Whitney U test, **Supplementary Fig. 9**). To avoid the correlation between length and other genomic features driving enrichment, we included length as a covariate in subsequent enrichment analyses.

To investigate the importance of regions overlapping misexpression-associated SVs versus controls, we tested for enrichment of five different genomic scores spanning evolutionary conservation, constraint and deleteriousness (**Fig. 3E, Supplementary Fig. 9, Methods**). Both misexpression-associated deletions and duplications were significantly enriched within more conserved regions compared to controls (Bonferroni p < 0.05) and were predicted to be significantly more deleterious by CADD-SV (Bonferroni p < 0.05)^21,22^. However, only duplications were located in more constrained regions (Bonferroni p < 0.05)^23,24^. Neither misexpression-associated deletions nor duplications were significantly enriched for human accelerated regions (HARs, Bonferroni p ≥ 0.05)^25^.

To determine whether misexpression-associated SVs were enriched in specific regulatory features compared to the control SVs, we annotated SVs with 23 regulatory features (**Fig. 3F, Supplementary Fig. 9, Methods**). Misexpression-associated deletions were most strongly enriched for transcribed regions but were also significantly enriched for regions with weak repressed polycomb, CTCF-binding sites in B-cells and neutrophils, CpG islands, enhancers and TAD boundaries (Bonferroni p < 0.05). Misexpression-associated duplications were most strongly enriched for enhancers but also showed significant enrichment for transcribed regions, active promoters, CpG islands and repressed polycomb (Bonferroni p < 0.05). Overall, these enrichments suggest that a subset of SVs may lead to gene misexpression via disruption of regulatory regions.

### Deletions and duplications lead to chimeric misexpression via transcriptional readthrough

Next, we aimed to identify putative mechanisms whereby the 105 misexpression-associated SVs lead to gene misexpression. From the genetic variant and regulatory feature enrichment analysis, we hypothesized that a subset of deletions and duplications could cause transcriptional readthrough resulting in chimeric gene misexpression. Based on their position and genomic context, we identified 17 (16.2%) transcriptional readthrough candidate SVs (12 deletions, 5 duplications) from the 105 misexpression-associated SVs (**Fig. 4A**, **Fig. 4B**, **Methods**).

**Figure 4.**
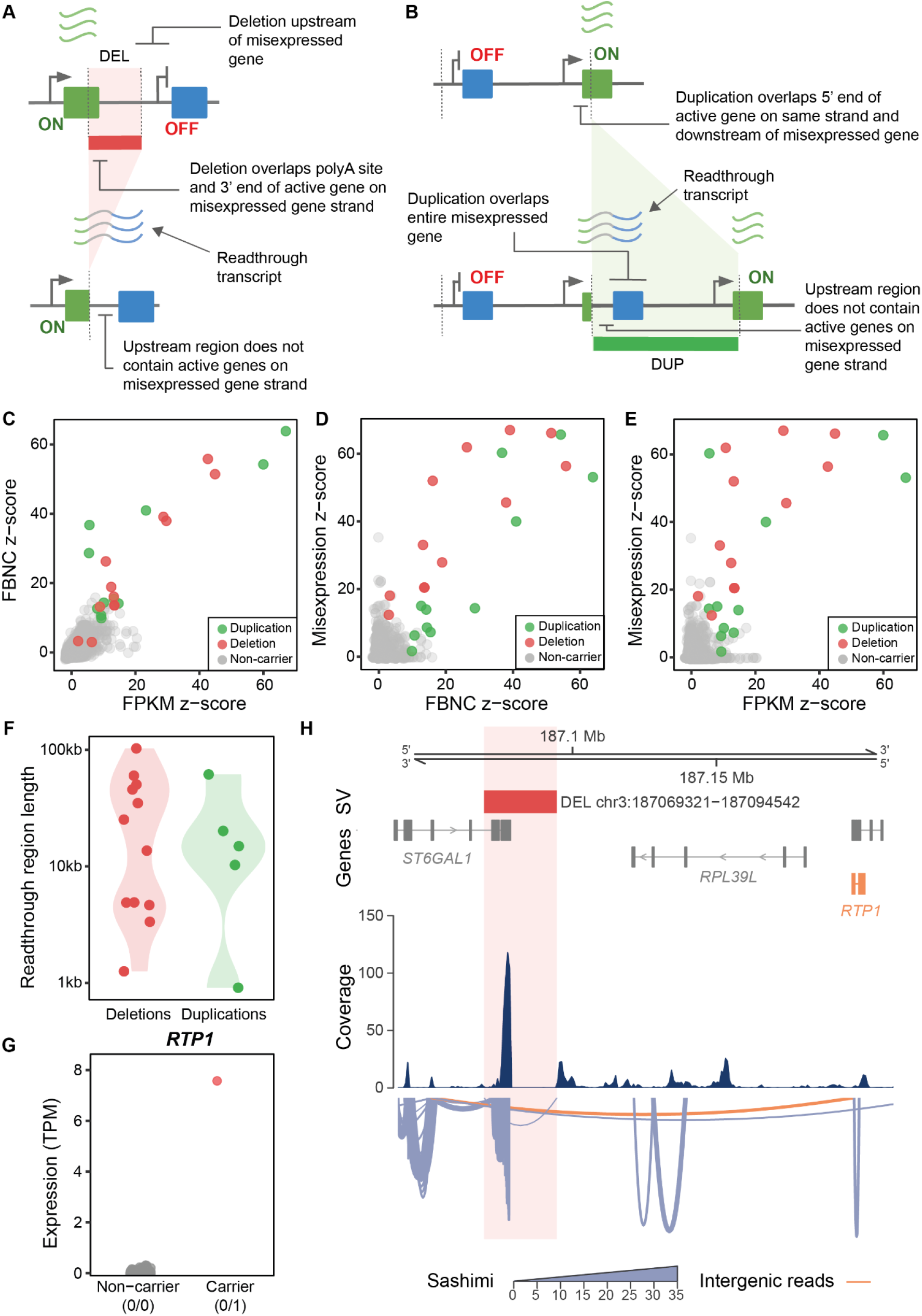
Transcriptional readthrough leads to chimeric misexpression. **A.)** Schematic diagram of a deletion resulting in transcriptional readthrough and chimeric gene misexpression. Deletion of the transcription termination site of an expressed gene (green) leads to transcriptional readthrough. This results in misexpression of the usually inactive gene (blue) located downstream. **B.)** Schematic diagram of a tandem duplication resulting in transcriptional readthrough and chimeric gene misexpression. The tandem duplication places an inactive gene (blue) downstream of an expressed gene (green) with no transcription termination site. This leads to transcriptional readthrough and misexpression of the usually inactive gene. **C.)** FPKM and FBNC z-scores over the predicted readthrough regions for carriers of candidate deletions (red) and duplications (green), as well as non-carriers (gray). Relationship between **D.)** FBNC z-score and **E.)** FPKM z-score with the respective misexpression z-score for carriers of candidate deletions (red) and duplications (green), as well as non-carriers (gray). **F.)** Length of the predicted readthrough region for duplications (red) and deletions (green). **G.)** Expression of RTP1 comparing a DEL chr3:187069321-187094542 carrier to non-carriers. Red color indicates samples passing the misexpression threshold TPM > 0.5 and z-score > 2 while gray samples are below this threshold. **H.)** Deletion of the 3’ end of ST6GAL1 results in transcriptional readthrough. Transcriptional readthrough leads to RTP1 misexpression (orange gene) and intergenic splicing between ST6GAL1 and RTP1 (intergenic reads, orange). In the sashimi plot, the line width corresponds to the number of reads spanning a given junction. FPKM; fragments per kilobase of transcript per million mapped reads, FBNC; fraction of bases with non-zero coverage. TPM; transcripts per million.

To assess whether these candidate SVs resulted in transcriptional readthrough we computed fragments per kilobase of transcript per million mapped reads (FPKM) and the fraction of bases with non-zero coverage (FBNC) over the predicted readthrough regions across all 4,568 RNA-seq samples (**Methods**). We z-score-transformed FPKM and FBNC metrics to account for differing levels of background transcription at each locus. For carriers of all 17 deletions and duplications, we observed aberrant (z-score > 2) levels of both FPKM and FBNC over the predicted readthrough region (**Fig. 4C, Supplementary Fig. 10**). Furthermore, both z-scores were positively correlated with the level of gene misexpression across carriers (FBNC Spearman’s rho = 0.76, p < 0.05, FPKM = 0.55, p < 0.05, **Fig. 4D**, **Fig. 4E**), while this was not the case for non-carriers (FBNC Spearman’s rho = 0.05 and FPKM = 0.09). Together these results provide strong evidence that these SVs lead to transcriptional readthrough resulting in gene misexpression.

Of the 12 transcriptional readthrough deletions, only 5 were within 5 kb and therefore were annotated by VEP as upstream variants with respect to the misexpressed gene (**Supplementary Fig. 10**). Of the 5 transcriptional readthrough duplications, 2 were annotated as non-coding transcript variants and 3 as transcript amplifications (**Supplementary Fig. 10**). These VEP consequences supported the enrichments observed in **Fig. 2E**. The median length of the readthrough region was 15 kb, but remarkably for one deletion (DEL chr3:187069321-187094542) we observed misexpression of a gene 103 kb away (**Fig. 4F**). At this locus, split reads revealed that intergenic splicing occurred between the expressed *ST6GAL1* gene and the usually inactive gene *RTP1* (**Fig. 4H**). The carrier of this deletion had highly aberrant expression levels of *RTP1* relative to non-carriers (**Fig. 4G**). *RTP1* is normally expressed in multiple non-blood tissues with the highest expression in the brain frontal cortex according to the GTEx project^26^. We also observed evidence of intergenic splicing due to transcriptional readthrough for a deletion (DEL chr5:77674588−77771600) involving misexpression of the *OTP* gene. According to the GTEx project, *OTP* is normally expressed in the hypothalamus (**Supplementary Fig. 10**, **Methods**)^26^.

### Deletions and duplications lead to chimeric misexpression via transcript fusion

Previous studies in rare diseases have demonstrated that pathogenic gene misexpression can occur via transcript fusion^7^. Therefore, we hypothesized that a subset of deletions and duplications could lead to chimeric gene misexpression via transcript fusion. To assess this, we used STAR fusion to identify fusion transcripts (**Methods**)^27^. We identified 12 fusion transcripts involving misexpressed genes that were consistently observed with a misexpression-associated SV within 200 kb. Out of these, we labeled 10 as high evidence and 2 as low evidence using STAR fusion’s filtering criteria (**Methods**), and focused our mechanistic analysis on fusion transcripts with high evidence. Of these, 3 were associated with duplications and 7 with deletions.

We had described 2 of the 7 deletion-associated fusion transcripts previously as being the result of transcriptional readthrough and intergenic splicing involving *RTP1* and *OTP* (**Fig. 4H**, **Supplementary Fig. 10**). Out of the other deletions, 2 removed the 3’ end and TTS of an active gene and 5’ end of an inactive gene on the same strand resulting in the inactive gene coming under the control of an active promoter (**Fig. 5A, Supplementary Fig. 11**). For the remaining 3 deletion-associated fusion events the mechanism was unclear. This may be due to failure to detect more complex rearrangements at these loci or because these variants are non-causal.

**Figure 5.**
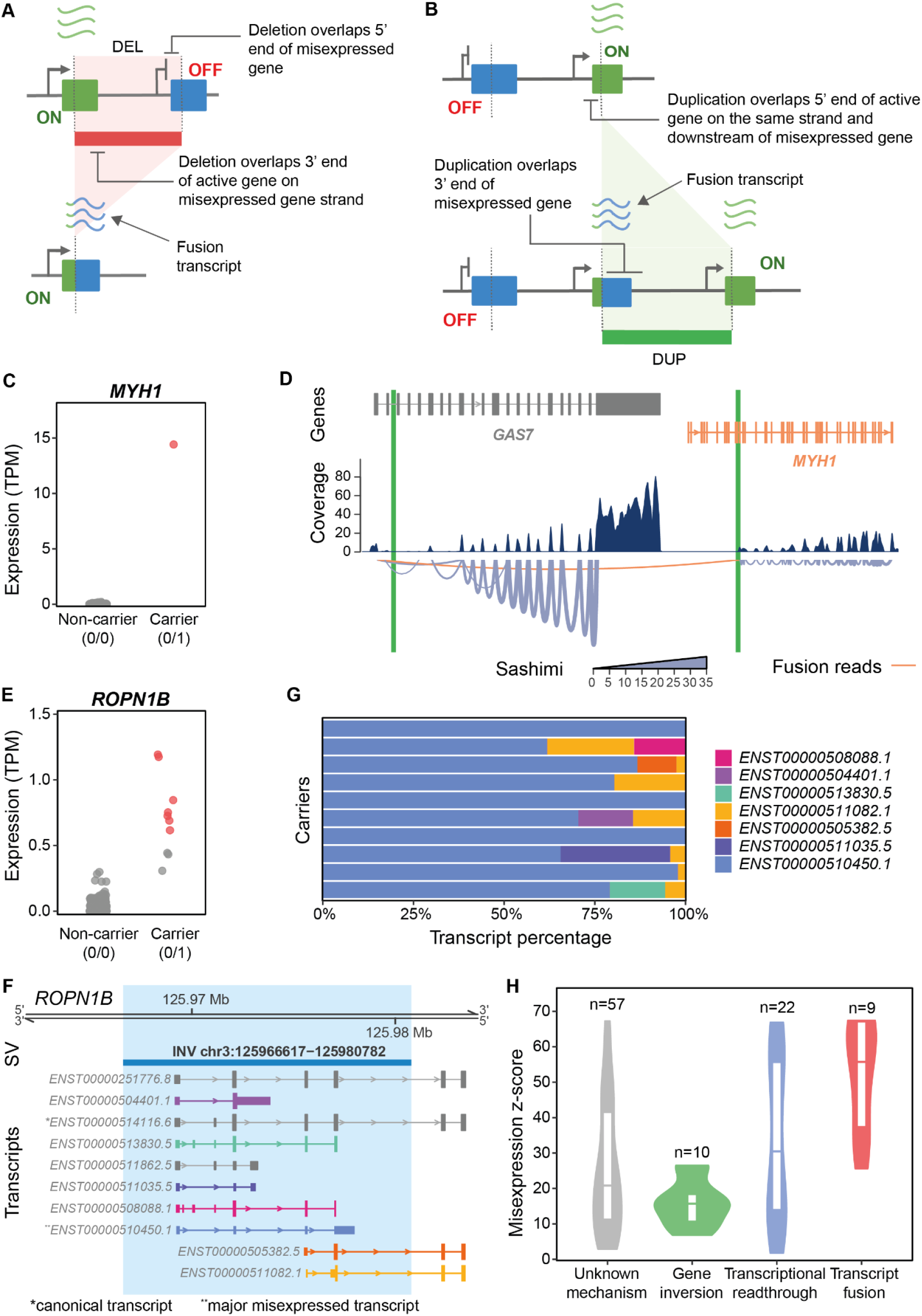
Transcriptional fusion and gene inversions lead to gene misexpression. ***A.)*** Schematic diagram of a deletion resulting in transcript fusion and chimeric gene misexpression. The deletion of the 3’ end of an active gene (green) and 5’ end of an inactive gene (blue) results in a fusion transcript containing portions of the active and inactive gene’s transcripts. **B.)** Schematic diagram of a tandem duplication resulting in transcript fusion and gene misexpression. The duplication of the 3’ end of an inactive gene (blue) and 5’ end of an active gene (green) in tandem results in a fusion transcript containing portions of the active and inactive gene’s transcripts. **C.)** Expression of MYH1 comparing a DUP chr17:10078018-10512685 carrier to non-carriers. Red color indicates samples passing the misexpression threshold TPM > 0.5 and z-score > 2 while gray samples are below this threshold. **D.)** FusionInspector visualization of the GAS7–MYH1 fusion transcript. DUP chr17:10078018-10512685 breakpoints are labeled in green. Introns have been shortened for visualization and breakpoint positions have been approximated accordingly. In the sashimi plot, the line width corresponds to the number of reads spanning a given junction. The misexpressed gene and the fusion reads are highlighted in orange. **E.)** Expression of ROPN1B comparing INV chr3:125966617-125980782 carriers and non-carriers. Red color indicates samples passing the misexpression threshold TPM > 0.5 and z-score > 2 while gray samples are below this threshold. **F.)** Location of INV chr3:125966617-125980782 showing all ROPN1B transcripts. The Ensembl canonical transcript is labeled with an asterisk and the major misexpressed transcript with a double asterisk. **G.)** Percentage expression of ROPN1B transcripts for all INV chr3:125966617-125980782 carriers. **H.)** Distribution of misexpression z-scores across different types of misexpression mechanisms. Text labels indicate the number of misexpression events for each putative mechanism.

All 3 duplications resulted in the 3’ end of an inactive gene being positioned within an active gene on the same strand (**Fig. 5B, Supplementary Fig. 11**). This leads to part of the inactive gene coming under the control of an active promoter. One of the duplications (DUP chr17:10078018-10512685) was associated with a *GAS7-MYH1* fusion transcript (**Fig. 5C**). The duplication’s breakpoints were consistent with the structure of the fusion transcript (**Fig. 5D**). *MYH1* is normally exclusively expressed in skeletal muscle tissue^26^. FusionInspector predicted that this fusion transcript contained a novel open reading frame resulting from the in-frame concatenation of 61 N-terminal residues of *GAS7* and 1603 C-terminal residues of *MYH1* (total predicted protein length 1664 residues)^28^. Since expression of this novel transcript is likely under the control of the *GAS7* promoter, we hypothesize that it will be misexpressed across multiple tissues^26^. This hypothesis is supported by the fact that fusion reads have been detected previously in multiple cancer samples from lung, stomach and intestine^29,30^. However, within our dataset this fusion transcript is likely the result of a heterozygous germline variant rather than a somatic variant.

### Inverting gene orientation is associated with non-chimeric misexpression

Gene misexpression was not limited to transcriptional readthrough and transcript fusion mechanisms. We observed that *ROPN1B* had consistently elevated expression across 10 carriers with an inversion (INV chr3:125966617-125980782) spanning the 5’ end of the gene (**Fig. 5E**, **Fig. 5F**). Transcript quantification using Salmon showed that *ROPN1B* misexpression was transcript-specific with the major misexpressed transcript being completely contained within the inversion (**Fig. 5G**)^31^. Inverting *ROPN1B’s* orientation may lead to ectopic enhancer-promoter contacts resulting in misexpression but this cannot be confirmed with the data available. Additionally, we observed elevated expression of intestinal alkaline phosphatase (*ALPI*) across 6 participants carrying two deletions (DEL chr2:232375546-232379537, and DEL chr2:232428106-232431877) in *cis* (**Supplementary Fig. 12**). However, the mechanism by which these deletions lead to misexpression is unclear.

Overall, we have identified a putative mechanism for 42% (41/98) of events with a misexpression-associated SV in *cis*. Out of these mechanisms, transcript fusion on average led to the most extreme levels of misexpression and gene inversions the lowest (**Fig. 5H**). We manually inspected the remaining 57 events but could not identify SVs with shared mechanisms that could explain the observed misexpression. Interestingly, only 4 of these events had an SV in *cis* that overlapped a TAD boundary and CTCF-binding site in the required orientation to result in misexpression via rearrangements in 3D chromatin architecture (**Methods**). However, we could not confirm that these SVs were causal. This result suggests that in our cohort 3D genome rearrangements leading to gene misexpression may be exceedingly rare.

## Discussion

In this study, we have developed gene misexpression as a novel type of transcriptomic outlier analysis and conducted the first genome-wide characterization of the gene misexpression landscape using bulk RNA-seq in a cohort of 4,568 blood donors. We found that misexpression events occur in the majority of samples and in over half of inactive genes. By integrating WGS and RNA-seq data, we assessed the influence of genetic variation on gene misexpression, demonstrating that these events are enriched for rare SVs in *cis*. We also show that a subset SVs lead to misexpression via specific mechanisms including transcriptional readthrough, transcript fusion and inverting gene orientation. These findings extend our understanding of gene misexpression and its genetic mechanisms beyond the limited number of samples and disease-relevant loci where misexpression had previously been described.

Large-scale RNA-seq studies have found that different categories of transcriptional outliers are enriched for distinct types of genetic variation^9,11^. We found that this was also the case for misexpression events, which were strongly enriched for rare SVs. Compared to SV enrichment in other outlier types measured across multiple tissues, this rare SV enrichment occurred at shorter distances from the TSS^11^. Although rare disease studies have demonstrated that SNVs and indels can lead to gene misexpression^6^, we did not observe a consistent significant enrichment for these types of genetic variation in this large predominantly healthy cohort. These results emphasize the disproportionate impact that large genetic perturbations have in influencing gene expression^9–12^. Unlike in previous studies focusing on SVs^10,12^, we identified multiple regulatory features that were enriched in misexpression-associated SVs and used these features to identify specific misexpression mechanisms.

Previous studies of rare diseases have observed that non-chimeric misexpression, where an individual inactive gene is aberrantly transcribed, can result from rearrangements in 3D chromatin architecture resulting in changes in enhancer-promoter interactions (enhancer adoption)^8^. However, in our cohort, we were unable to identify such events with confidence. This might indicate that these events are exceedingly rare in healthy human populations. Alternatively, our approach may fail to identify these events due to a lack of high-resolution, context-specific Hi-C data. However, parallels can be drawn between the SV mechanisms resulting in chimeric misexpression we have observed here and those involving rearrangements in 3D chromatin architecture. While SVs leading to chimeric misexpression via transcript fusion or transcriptional readthrough place an inactive gene under the control of a different active promoter, SVs resulting in enhancer adoption place an inactive promoter under the control of active enhancers. Therefore, a consistent theme across different SV misexpression mechanisms is changes to the regulatory environment of an inactive gene through alterations to either its promoter or enhancers.

It is important to highlight the limitations of this study. Firstly, we have only analyzed gene misexpression within whole blood and further studies should examine the prevalence of misexpression across different tissues and cell types as well as in disease. Indeed, a link between misexpression and disease is more likely to be detected in the relevant disease tissue rather than in whole blood. Secondly, we have focused on high confidence events by using a stringent expression threshold. This threshold is arbitrary and the level of misexpression required to influence cellular processes is likely to vary across genes and contexts. Thirdly, due to the technical difficulties of calling SVs in short-read genome sequencing, we may be unable to detect some SVs that lead to misexpression and therefore the proportion of misexpression events associated with an SV is likely underestimated. Some of the misexpression events not associated with SVs may be due to non-genetic mechanisms such as leaky transcription, chromatin plasticity or specific environmental cues. Finally, when interpreting the consequences of rare SVs we have focused on mechanisms that are shared by multiple variants or events with multiple variant carriers. Therefore, we are biased towards detecting more common misexpression mechanisms and may miss additional mechanisms caused by ultra-rare SVs.

Interpreting the functional effects of rare genetic variation remains challenging and is important for understanding the molecular mechanisms by which variants influence human traits. Here, we have extended our understanding of how genetic variants influence gene expression. The fact that rare SVs can induce misexpression not just in the rare disease context should be taken into account in future studies when cataloging and interpreting their effects in population cohorts. This is especially important for SVs associated with human complex diseases, as it is currently unknown what fraction of these SVs may mediate their phenotypic effects by causing gene misexpression.

## Methods

### The INTERVAL Study

The INTERVAL study is a prospective cohort study of approximately 50,000 participants nested within a randomized trial of varying blood donation intervals^14,15^. Between 2012 and 2014, blood donors aged 18 years and older were recruited at 25 centers of England’s National Health Service Blood and Transplant (NHSBT). All participants gave informed consent before joining the study and the National Research Ethics Service approved this study (11/EE/0538). Participants were generally in good health as blood donation criteria exclude individuals with a history of major diseases (e.g. myocardial infarction, stroke, cancer, HIV, and hepatitis B or C) and who have had a recent illness or infection. Participants completed an online questionnaire comprising questions on demographic characteristics (e.g. age, sex, ethnicity), lifestyle (e.g. alcohol and tobacco consumption), self-reported height and weight, diet and use of medications.

### Whole genome sequencing

The manuscript describing the whole genome sequencing in full is in preparation. In brief, whole genome sequencing was performed on 12,354 samples using the Illumina HiSeq X10 platform as paired-end 151 bp reads at the Wellcome Sanger Institute (WSI). Reads were aligned to the GRCh38 human reference genome with decoys (also known as HS38DH) using BWA MEM. Variants were called using GATK4.0.10.1. GATK Variant Quality Score Recalibration (VQSR) was used to identify probable false positive calls. 491 samples were removed including 77 samples with coverage below 12x, 134 with > 3% non-reference discordance (NRD), 118 with > 3% FreeMix (VerifyBamID2) score, 221 samples failing identity checks, 30 samples swapped, 40 samples failing sex checks, 39 duplicates and 9 samples with possible contamination. Genotypes with allele read balance > 0.1 for homozygous reference variants, < 0.9 for homozygous alternative variants or not between 0.2-0.8 for heterozygous variants were removed. Genotypes were also removed if the proportion of informative reads was < 0.9 or total read depth > 100. We performed variant quality control and filtered out variants that failed to meet the following requirements: call rate per site > 95%, mean genotype quality (GQ) value > 20, Hardy-Weinberg equilibrium (HWE) p-value > 1 x 10^-6^ only for autosomes. All monomorphic variants with alternative allele count (AAC) = 0 were further removed, although we kept all monomorphic variants with reference allele count (RAC) = 0. For chrX and chrY we applied an additional step to correct the allele counts and frequencies due to female and male samples. Overall this resulted in 116,382,870 variants (100,694,832 SNVs and 15,688,038 indels) including 6,637,420 (5.7%) multi-allelic sites across 11,863 participants.

### Structural variant calling

Generation of the SV callset has been described in full previously^32^. In brief, deletions, duplications, inversions and mobile element insertions were called using a combination of Genome STRiP^33^, Lumpy^34^, CNVnator^35^ and svtools^36^. For duplications and deletions, a random forest classifier using read alignment parameters was trained to minimize false positives. This resulted in 88% sensitivity and 99% specificity for deletions, and 55% sensitivity and 92% specificity for duplications. Inversions were retained if < 10% of genotypes were missing, HWE was not violated, and < 10% of alternate allele supporting reads came from split and paired read ends. Breakends were removed from the callset. Final tuning of the overall quality score was modeled to ensure that 90% of carrier genotypes were identical among duplicate samples. SVs with significant overlap across carriers were collapsed using a graph-based procedure. The final callset consisted of 123,801 SVs comprising 107,966 deletions, 11,681 duplications, 1,395 inversions and 1,395 mobile element insertions across 10,728 participants. The final callset was compared to SV calls from the 1000 genomes and Hall-SV cohorts. The callset captured 93% and 92% of common deletions and 65% and 75% of common duplications, from each cohort respectively.

### RNA sample processing and sequencing

The manuscript describing the RNA sequencing data in full is in preparation. In brief, blood samples were collected from INTERVAL participants in Tempus Blood RNA Tubes (ThermoFisher Scientific) and stored at -80°C until use. RNA extraction was performed by QIAGEN Genomic Services using an in-house developed protocol. mRNA was isolated using a NEBNext Poly(A) mRNA Magnetic Isolation Module (NEB) and then re-suspended in nuclease-free water. Globin depletion was performed using a KAPA RiboErase Globin Kit (Roche). RNA library preparation was done using a NEBNext Ultra II DNA Library Prep Kit for Illumina (NEB) on a Bravo WS automation system (Agilent). Samples were PCR amplified using a KapaHiFi HotStart ReadyMix (Roche) and unique dual-indexed tag barcodes. PCR products were purified using AMPure XP SPRI beads (Agencourt). Libraries were pooled up to 95-plex in equimolar amounts on a Biomek NX-8 liquid handling platform (Beckman Coulter), quantified using a High Sensitivity DNA Kit on a 2100 Bioanalyzer (Agilent), and then normalized to 2.8 nM. Samples were sequenced using 75 bp paired-end sequencing (reverse stranded) on a NovaSeq 6000 system (S4 flow cell, Xp workflow; Illumina).

### RNA sequencing alignment

The data pre-processing, including RNA-seq quality control, STAR and Salmon alignments were performed with a Nextflow pipeline, which is publicly available at https://github.com/wtsi-hgi/nextflow-pipelines/blob/rna_seq_interval_5591/pipelines/rna_seq.nf, including the specific aligner parameters. We assessed the sequence data quality using FastQC v0.11.8. Reads were aligned using STAR v2.7.3.a^37^. The STAR index was built against GRCh38 Ensembl GTF v97 using the option -sjdbOverhang 75. STAR was run in a two-pass setup with recommended ENCODE options to increase mapping accuracy: (i) a first alignment step of all samples was used to discover novel splice junctions; (ii) splice junctions of all samples from the first step were collected and merged into a single list; (iii) a second step realigned all samples using the merged splice junctions list as input. From the aligned RNA-seq read data, gene-level read counts were calculated from the number of reads mapping to exons using featureCounts v2.0.0^38^. The raw gene-level count data contained 60,617 genes across 4,778 samples.

### Quality control of RNA sequencing samples

Samples mismatched between RNA-seq and genotyping data within the cohort were identified using QTLtools MBV v1.2^39^. Five sample swaps were corrected. Samples with covariates indicating lower quality data were identified and removed using the following metrics: RIN < 4 or read depth < 10 million assigned reads by featureCounts v2.0.0^38^. Samples with missing sequencing covariates and genotyping data were removed as well as samples with suspected contamination. One sample from each flagged pair of related participants, estimated as first- or second-degree from genetic data, was removed, prioritizing samples with WGS data. After this stage, 47 samples were removed, leaving 4,731 remaining samples.

### Gene expression quantification

Prior to expression quantification the following genes were removed: globin genes, rRNA genes, genes on non-reference chromosomes, pseudoautosomal region genes and genes with transcripts annotated as “retained_intron” or “read_through”. After removing these genes, 59,144 remained. Gene-level read counts were converted to TPM values using the total length of merged exons. The total length of merged exons was computed by collapsing the GENCODE Release 31 annotation to a single transcript model for each gene using the custom isoform collapsing procedure from the GTEx project^26,40^.

### Removal of global expression outliers

Using TPM values across 59,144 genes and 4,731 samples, samples with many top expression events due to either technical or biological effects were removed as global expression outliers. To count the number of top expression events per sample, genes with TPM equal to zero across all samples were not included leaving 57,555 genes. Then, for each sample the number of most extreme expression events in these remaining genes was calculated. 3.4% of samples (163/4,731) with ≥5x the expected number of top expression outliers (total genes/total samples) were removed resulting in 4,568 samples (**Supplementary Fig. 1**).

### Inactive gene identification

Enrichment testing and downstream analysis were limited to autosomal protein-coding or long non-coding RNA genes in GENCODE Release v31^40^. Additionally, to minimize misexpression false positives only genes that passed the expression thresholds for expression quantitative trait loci (eQTL) mapping in at least one of 49 GTEx tissues were retained, leaving 29,614 genes^26^. For the remaining genes, we calculated the percentage of samples with a TPM > 0.1. Across all genes, this percentage had a bimodal distribution separating highly and lowly expressed genes (**Fig. 1B**). To focus on genes that had very low or no detectable expression, we selected 8,779 genes which had a TPM > 0.1 in less than 5% of samples. This approach is analogous to the method used by the GTEx consortium to define active genes for eQTL mapping^26^.

### Inactive gene set validation

To validate our inactive gene set we used several approaches:

1. We intersected 60,603 genes in GENCODE Release v31 with predicted chromatin states from the Roadmap Epigenomics Project’s 15-state ChromHMM trained on PBMC data^41^. Then, for each gene we calculated the fractional overlap of each chromatin state and conducted k-means clustering to group genes that had similar epigenetic modifications. We performed k-means clustering with 2-10 clusters and selected k = 8 clusters because these clusters were the most biologically interpretable. Based on the overlapping chromatin states, we manually annotated these 8 clusters with the following labels: transcription, weak transcription, quiescent, polycomb weak quiescent, polycomb weak, polycomb repressed, heterochromatin, and unassigned (**Supplementary Fig. 1**). 94% (8,277/8,779) of inactive genes were grouped in clusters with high overlap of repressive or quiescent chromatin states (**Supplementary Fig. 1**).
2. We checked whether our method of identifying inactive genes led to a similar gene set using a different whole blood RNA sequencing dataset. We identified inactive genes in whole blood RNA sequencing data from GTEx using the same approach. In brief, we focused on 558 individuals with European ancestry to limit the effects of population stratification. Samples with ≥5x the expected number of top expression outliers (total genes/total samples) were removed (n = 18). Comparison between INTERVAL and GTEx was restricted to the 29,614 genes defined previously. Inactive genes were defined as having a TPM > 0.1 in less than 5% of samples (n = 8,207). 80% of the INTERVAL inactive gene set were found in the inactive genes identified in GTEx. This is in spite of cohort differences including sample size, participant age, transcript annotation reference, RNA-seq strandedness and sampling method (blood donation versus post-mortem).
3. We tested whether our inactive gene set contained many genes expressed in other whole blood RNA sequencing datasets. To do so, we examined the overlap between inactive genes and eGenes (genes with at least one eQTL) from GTEx and eQTLgen^26,42^. Only 6.4% and 3.1% of the INTERVAL inactive genes were eGenes in GTEx whole blood and eQTLGen, respectively.

These results confirmed that we had identified a set of genes with very low or no expression across different datasets using information from different data types.

### Defining gene misexpression

TPM values were z-score transformed for each inactive gene across all 4,568 samples passing quality control. A gene in a sample was defined as misexpressed if it had a TPM > 0.5 and a z-score > 2. Z-scores were used to allow comparison of misexpression events across genes. In addition to the z-score threshold, a TPM threshold of 0.5 was used to remove misexpression events that had a high z-score but only low expression.

### Accounting for non-genetic sources of gene misexpression

To ensure that gene misexpression was not associated with biological or technical confounders we correlated the expression of each inactive gene with 225 covariates. These covariates included sample age, height, weight, BMI, sex, 89 Sysmex cell count measurements, 67 inferred xCell cell enrichments^43^, 25 technical covariates, top 20 genetic PCs, season and sequencing batch. We removed 1.5% (129/8779 genes) whose expression was significantly correlated (|Spearman’s rho| > 0.2, FDR-adjusted p < 0.05) with any covariate (**Supplementary Table 1**). The low percentage of genes removed confirmed that for the majority of genes, misexpression events could not be attributed to the systematic effect of a measured or inferred covariate. The final inactive gene set is provided in **Supplementary Table 2**.

### Gene-level features

We curated a set of gene-level features in order to understand the differences between misexpressed and non-misexpressed genes. The full set of features is provided in **Supplementary Table 3**. For each of the 8,650 inactive genes we compiled a set of 81 gene-level features across 5 major categories: genomic, constraint and conservation, expression, regulation and gene sets. Genomic features such as gene length, gene density and distance to the closest gene were calculated from GENCODE Release v31^40^. Constraint scores included LOEUF and missense OUEF from gnomAD, as well as pLI, probability of recessive lethality (pRec), probability of complete haploinsufficiency (pNull), pHaplo, pTriplo, episcore and the enhancer domain score (EDS)^16–18,44^. The mean conservation score across the gene body was calculated using PhyloP basewise conservation score across 100 vertebrates^21^. The number of conserved elements per base pair within a ±10 kb around a gene was calculated using GERP++ conserved elements^45^. The number and type of tissues a gene is expressed in was calculated from GTEx^26^. Active gene density and distance were calculated by subsetting to genes with a median TPM > 0.5 in INTERVAL. Regulation features derived from chromatin states were calculated using the Roadmap Epigenomics Project’s 15-state ChromHMM trained on PBMC data^41^. The fraction of a gene overlapping A/B compartments was derived from GM12878 Hi-C data processed by the 4D Nucleome Program^46,47^. We generated a set of TAD boundaries in GM12878 cells that were shared (within ±50 kb) across IMR90, HUVEC, HNEK and HMEC cell lines from the 4D Nucleome Program and used these shared boundaries to calculate the closest distance from each gene to a TAD boundary. Enhancer features based on proximity- and activity-linking methods were from Wang & Goldstein^17^. Gene sets included oncogenes (tier 1, dominant) from the COSMIC v97 Cancer Gene Census (CGC)^48^, approved drug targets curated by OpenTargets (OT release v22.11)^49^, developmental disorder genes from the Decipher DDG2P database^50^ and OMIM^19^, and different gene sets annotated by gnomAD including olfactory, autosomal recessive, autosomal dominant and haploinsufficient genes^16^. All features were z-score transformed across all inactive genes. Inactive genes were split into two groups depending on whether they were not misexpressed (4,437 genes) or misexpressed at least once (4,213 genes) defining misexpression with a misexpression z-score > 2 and TPM > 0.5. Using different z-score thresholds did not lead to markedly different results (**Supplementary Fig. 3**). The enrichment of each feature within the misexpressed group was calculated using logistic regression. Across all tests, p-values were adjusted using Bonferroni correction. 95% confidence intervals for the fitted parameters were calculated using the standard normal distribution. Underenrichment of HPO terms within misexpressed genes was calculated using the gProfiler (gProfiler2 v0.2.1) functional profiling function with all 8,650 inactive genes used as the custom background^51^.

### Matching RNA sequencing and whole genome sequencing samples

Samples with matching RNA-seq and WGS data were identified using QTLtools MBV v1.2^39^. Out of 4,568 RNA-sequencing samples, 2,821 and 2,640 samples had a matching WGS sample with SNV/indel calls and SV calls, respectively. The difference in matching samples was due to a higher number of samples failing SV calling.

### Genetic variant enrichment calculations

For each enrichment test we defined a misexpression group as all expression events (expression of a given gene in an individual) passing the specified misexpression z-score threshold and a TPM > 0.5. The control group was defined as all expression events below these thresholds restricted to the genes within the misexpression group. Therefore, for each misexpression threshold the misexpression and control group gene sets were identical. This ensured that enrichment calculations reflected differences in genetic effects rather than differences in mutation background distributions between non-identical gene sets. Risk ratios were calculated as the proportion of expression events in the misexpression group with a given variant type within the tested genomic region and MAF range over the proportion of events in the control group. For SV enrichment tests we counted variants overlapping a ±200 kb window around the gene body while for SNVs and indels a ±10 kb window was used. P-values were calculated using a two-sided Fisher’s exact test and 95% confidence intervals were calculated using a normal approximation. We tested four non-overlapping MAF thresholds: rare (0-1% MAF), low-frequency (1-5% MAF) and common variants (5-10% and 10-50% MAF).

To test the enrichment of variants at different genomic distances from genes involved in misexpression events, we assigned variants to a genomic window for each gene. Variants were assigned to 200 kb windows up to 1 Mb upstream and downstream from the gene start and end, respectively, or when overlapping the gene itself to the gene body window. In cases where a variant spanned multiple windows, the variant was placed in the window closest to the gene with variants overlapping any part of the gene assigned to the gene body window. This resulted in all gene-variants pairs being uniquely assigned to a single window. Enrichment testing was then conducted for each genomic window separately.

To investigate variant consequences, SVs were annotated with the most severe VEP consequence on the gene in the test window (VEP v97.3)^20^. Variants with no predicted consequence on the gene were annotated according to the most severe consequence if the annotation was regulatory or intergenic (TFBS ablation, TF_binding_site_variant, regulatory_region_variant, TFBS_amplification, intergenic_variant, regulatory_region_ablation, regulatory_region_amplification). If the variant had no predicted consequence on the gene and its most severe consequence was not regulatory or intergenic then it was annotated as having no predicted effect. Enrichment calculations were performed for each variant consequence that had at least one carrier within the tested window. For SVs a ±200 kb window around the gene body was used.

Overall, we performed 700 genetic variant enrichment tests. Across all tests, p-values were adjusted using Bonferroni correction. All generic variant enrichment results can be found in **Supplementary Table 4.**

### Identifying misexpression-associated and control rare structural variants

We identified 23,159 rare (MAF < 1%) SVs located within ±200 kb of an inactive gene for which misexpression (z-score > 2 and TPM > 0.5) was observed at least once (4,437 genes). For each gene-SV pair, we calculated the median TPM and z-score across all carriers. We defined misexpression-associated SVs as SVs with a nearby gene that had a median TPM > 0.5 and median z-score > 2. We additionally excluded gene-SV pairs where any carrier had a TPM < 0.1, resulting in 105 misexpression-associated SVs. These criteria allow for variable levels of gene expression around the misexpression threshold while removing likely non-causal variants. From the 23,159 rare SVs, we defined control SVs as having a maximum TPM equal to 0 for every inactive gene where the SV is within 200 kb. This resulted in 20,157 control variants.

Misexpression-associated SVs were annotated based on their VEP consequence on the misexpressed gene as done for the genetic enrichment analysis and their position relative to the misexpressed gene. We confirmed that all misexpression-associated duplications were tandem duplications by manually inspecting them in the Integrative Genome Viewer^52^.

### Structural variant properties

To understand the different properties of misexpression-associated and control SVs we annotated SVs with 5 features based on conservation, mutational constraint and deleteriousness scores. Deletions and duplications were scored with CADD-SV v1.1 in batches of 5000 variants^22^. Scoring inversions and mobile element insertions is currently not supported by CADD-SV. Since we were comparing CADD-SV distributions between control and misexpression-associated variants we used the raw CADD-SV scores as recommended by the authors. PhyloP conservation scores were downloaded from UCSC genome browser^21^. Each SV was annotated based on the maximum conservation score observed across all overlapping bases. Constraint z-scores passing all quality control checks for coding and non-coding regions were downloaded from gnomAD^24^. SVs were annotated with the maximum gnomAD z-score across all overlapping 1 kb windows. SVs were annotated based on the minimum gwRVIS across all overlapping bases^23^. SVs were annotated with a categorical variable based on whether they overlapped a HAR^25^. Misexpression-associated deletions and duplications were compared separately versus controls using logistic regression with each score modeled independently, z-score transformed and SV length included as a covariate. P-values were adjusted across all tests using Bonferroni correction.

95% confidence intervals for the fitted parameters were calculated using the standard normal distribution. Excluding SV length as a covariate did not lead to dramatic changes in the enrichments (**Supplementary Fig. 8**).

### Structural variant regulatory features

To understand the regulatory features specific to misexpression-associated SVs, we conducted enrichment analysis across 23 regulatory features. All regulatory features with their transformation, cell type and datasource are described in **Supplementary Table 5**. A/B compartments measured in the GM12878 cell-line were downloaded from the 4D nucleome project^46,47^. We generated a set of TAD boundaries in GM12878 cells that were shared across IMR90, HUVEC, HNEK and HMEC cell lines. A TAD boundary was considered shared if another TAD boundary was located within ±50 kb in another cell line. CpG islands were downloaded from the UCSC genome browser^53^. CTCF-only candidate cis-regulatory elements (cCREs) across all cell types were downloaded from ENCODE^54^. CTCF cCREs generated in primary cells from whole blood with CTCF ChIP-seq data available (CD14+ monocytes, neutrophils, and B-cells) were also downloaded^54^. Regulatory features derived from chromatin states were calculated using the Roadmap Epigenomics Project’s 15-state ChromHMM trained on PBMC data^41^. For all data types, features were generated by encoding SV overlap as a binary indicator that was subsequently z-score transformed.

To assess the enrichment of different regulatory annotations in misexpression-associated SVs, we performed logistic regression modeling misexpression status as a function of each regulatory feature individually with SV length included as an additional covariate. Excluding SV length did not lead to dramatic changes in the log-odds values but many more features were significant (**Supplementary Fig. 8**). Logistic regression was conducted separately for deletions and duplications. P-values were adjusted across all tests using Bonferroni correction. 95% confidence intervals for the fitted parameters were calculated using the standard normal distribution.

### Selection and characterisation of transcriptional readthrough candidate structural variants

To identify deletions that were transcriptional readthrough candidates, we selected misexpression-associated deletions that satisfied the following criteria:

1. The deletion is located upstream of the misexpressed gene.
2. The deletion partially overlaps a gene’s 3’ end and overlaps a terminal exon polyA site from polyASite 2.0^55^. The overlapping gene is expressed in whole blood (median TPM > 0.5 in the INTERVAL dataset) and is on the same strand as the misexpressed gene.
3. The region upstream of the misexpressed gene up to the SV breakpoint does not contain an expressed gene (median TPM > 0.5 in the INTERVAL dataset) on the same strand as the misexpressed gene.

If a deletion was associated with misexpression of multiple genes then the gene closest to the SV was selected in order to define the expected readthrough region.

To identify duplications that were transcriptional readthrough candidates, we selected misexpression-associated duplications that satisfied the following criteria:

1. The duplication overlaps the entire misexpressed gene.
2. The duplication partially overlaps the 5’ end of a gene that is expressed (median TPM > 0.5 in the INTERVAL dataset), is positioned downstream of the misexpressed gene and is on the same strand as the misexpressed gene.
3. The region upstream of the misexpressed gene up to the SV breakpoint does not contain an expressed gene (median TPM > 0.5 in the INTERVAL dataset) on the same strand as the misexpressed gene.

If a duplication was associated with misexpression of multiple genes then the gene with the shortest expected readthrough region was selected. This resulted in 12 transcriptional readthrough candidate deletions and 5 transcriptional readthrough candidate duplications.

For both deletions and duplications, the read count and fraction of bases with non-zero coverage of the region upstream of the misexpressed gene up to the SV breakpoint was calculated using BedTools coverage requiring the same strandedness and treating split BAM entries as distinct bed intervals. Read counts were converted to FPKM using sample read depth and the length of the region. For each readthrough region, z-scores were calculated for both FPKM and FBNC metrics across all 4568 RNA-seq samples passing quality control. 2640 samples with available SV calls and WGS were then annotated as either deletion carriers, duplication carriers or non-carriers.

### Identification of fusion transcripts

We used STAR fusion v.1.10.1 to identify fusion transcripts^27^. First, we ran STAR fusion across all samples with a misexpression-associated variant in max sensitivity mode with a STAR max mate distance of 50 kb and with no annotation filter (as recommended for detecting fusion in non-cancer samples). We selected fusion events that involved misexpressed genes that had a misexpression-associated SV in *cis*. We removed fusion events that were not supported across all carriers of the misexpression-associated SV. Next, we ran STAR fusion again using the same parameters except without applying the max sensitivity mode and with the –denovo_reconstruct, FusionInspector validate and –examine_coding_effect flags applied^28^. Fusion transcripts validated by FusionInspector from this run were labeled high evidence while those that were only identified in the first run were labeled low evidence.

### Salmon transcript quantification

We used Salmon v1.1.0 for transcript quantification^31^. The Salmon index was built against GRCh38 cDNA, which was used to generate transcript level quantification from the sequence data. R packages tximport v1.14.2, AnnotationHub v2.18.0, BiocFileCache v1.10.2, BiocGenerics v0.32.0 were applied to obtain various count matrices from these quantifications at the transcript or gene level. For INV chr3:125966617-125980782 carriers the transcript percentage for each *ROPN1B* transcript was calculated as the transcript TPM divided by the total TPM across all transcripts.

### Identification of SVs with potential to alter 3D chromatin architecture

From the misexpression events with no putative mechanism we selected misexpression-associated SVs that overlapped both a TAD boundary (shared across multiple cell-lines), a CTCF-only cCRE across all cell-types from ENCODE^46^. For duplications, we also required that the variant completely overlapped the misexpressed gene and an Enh or EnhG chromHMM state from PBMCs^41^. For deletions, we also required that the variant did not overlap the misexpressed gene. This led to the identification of 4 misexpression events with a candidate SV for altering 3D chromatin architecture.

### Genome track visualization

Gviz v.1.38.4 was used to visualize genomic tracks and FusionInspector results^56^.

## Data availability

The INTERVAL study data used in this paper are available to bona fide researchers from. The data access policy for the data is available on emailing. The RNA-seq data (n = 4,732 INTERVAL participants) have been deposited at the European Genome-phenome Archive (EGA) under the accession number EGAD00001008015. The WGS data have been deposited at the EGA under accession number EGAD0000100866.

GENCODE v31: https://ftp.ebi.ac.uk/pub/databases/gencode/Gencode_human/release_31/gencode.v31.annotation.gtf.gz

Ensembl v97: http://ftp.ensembl.org/pub/release-97/gtf/homo_sapiens/Homo_sapiens.GRCh38.97.gtf.gz

eQTLGen eQTL: https://molgenis26.gcc.rug.nl/downloads/eqtlgen/cis-eqtl/2019-12-11-cis-eQTLsFDR-ProbeLevel-CohortInfoRemoved-BonferroniAdded.txt.gz

GTEx v8 eQTL expression matrices: https://storage.googleapis.com/gtex_analysis_v8/single_tissue_qtl_data/GTEx_Analysis_v8_eQTL_expression_matrices.tar

GTEx v8 eQTL data all tissues: https://storage.googleapis.com/gtex_analysis_v8/single_tissue_qtl_data/GTEx_Analysis_v8_eQTL_EUR.tar

GTEx v8 read count whole blood: https://storage.googleapis.com/gtex_analysis_v8/rna_seq_data/gene_tpm/gene_tpm_2017-06-05_v8_whole_blood.gct.gz

GTEx v8 median TPM per tissue: https://storage.googleapis.com/gtex_analysis_v8/rna_seq_data/GTEx_Analysis_2017-06-05_v8_RNASeQCv1.1.9_gene_median_tpm.gct.gz

OMIM: https://www.omim.org

DECIPHER gene list: https://www.ebi.ac.uk/gene2phenotype/downloads/DDG2P.csv.gz

pHaplo and pTriplo scores: https://ars.els-cdn.com/content/image/1-s2.0-S0092867422007887-mmc7.xlsx

ChromHMM PBMC states: https://egg2.wustl.edu/roadmap/data/byFileType/chromhmmSegmentations/ChmmModels/coreMarks/jointModel/final/E062_15_coreMarks_hg38lift_mnemonics.bed.gz

ENCODE cCREs all CTCF-only sites: https://downloads.wenglab.org/Registry-V3/GRCh38-cCREs.CTCF-only.bed

ENCODE cCREs CD14+ monocytes: https://downloads.wenglab.org/Registry-V3/Seven-Group/ENCFF389PZY_ENCFF587XGD_ENCFF184NWF_ENCFF496PSJ.7group.bed

ENCODE cCREs B-cells: https://downloads.wenglab.org/Registry-V3/Seven-Group/ENCFF035DJL.7group.bed

ENCODE cCREs Neutrophils: https://downloads.wenglab.org/Registry-V3/Seven-Group/ENCFF685DZI_ENCFF311TAY_ENCFF300LXQ.7group.bed

CpG islands: http://hgdownload.soe.ucsc.edu/goldenPath/hg38/database/cpgIslandExt.txt.gz

gnomAD metrics: https://storage.googleapis.com/gcp-public-data--gnomad/release/2.1.1/constraint/gnomad.v2.1.1.lof_metrics.by_transcript.txt.bgz

gnomAD constraint z-scores: https://gnomad.broadinstitute.org/downloads#v3-genomic-constraint

gnomAD gene lists: https://static-content.springer.com/esm/art%3A10.1038%2Fs41586-020-2308-7/MediaObjects/41586_2020_2308_MOESM4_ESM.zip

gwRVIS: https://az.app.box.com/v/jarvis-gwrvis-scores/folder/159704875574

GERP++ elements: https://bds.mpi-cbg.de/hillerlab/120MammalAlignment/Human120way/data/conservation/gerpElements_hg38_multiz120Mammals.bed.gz

PhyloP 100-way: http://hgdownload.soe.ucsc.edu/goldenPath/hg38/phyloP100way/hg38.phyloP100way.bw

HARs: https://ftp.ncbi.nlm.nih.gov/geo/series/GSE180nnn/GSE180714/suppl/GSE180714%5FHARs%2Ebed%2Egz

Cosmic Cancer Gene Census v97: https://cancer.sanger.ac.uk/census

OpenTargets targets information: http://ftp.ebi.ac.uk/pub/databases/opentargets/platform/22.11/output/etl/json/targets/

A/B compartments in GM12878: https://data.4dnucleome.org/files-processed/4DNFILYQ1PAY/

TAD boundaries in GM12878: https://data.4dnucleome.org/files-processed/4DNFIVK5JOFU/

TAD boundaries in HMEC: https://data.4dnucleome.org/files-processed/4DNFIJL18YS3/

TAD boundaries in HNEK: https://data.4dnucleome.org/files-processed/4DNFICLU9GUP/

TAD boundaries in HUVEC: https://data.4dnucleome.org/files-processed/4DNFI9MZWZF7/

TAD boundaries in IMR90: https://data.4dnucleome.org/files-processed/4DNFIMNT2VYL/

polyASite 2.0 database polyA sites: https://polyasite.unibas.ch/download/atlas/2.0/GRCh38.96/atlas.clusters.2.0.GRCh38.96.bed.gz

polyASite 2.0 database polyA sites with sample information: https://polyasite.unibas.ch/download/atlas/2.0/GRCh38.96/atlas.clusters.2.0.GRCh38.96.tsv.gz

## Code availability

The Nextflow pipeline used for STAR and Salmon alignments is available at https://github.com/wtsi-hgi/nextflow-pipelines/blob/rna_seq_interval_5591/pipelines/rna_seq.nf. Custom code used for analysis of processed sequencing data is available here: https://github.com/tvdStichele/interval_misexpression_manuscript.

## Author contributions

Conceptualisation: T.V.; data analysis: T.V., K.L.B., W.L., B.H., K.W., K.K., E.P., J.M., A.P.N.; provided resources: B.H., K.W., K.K., E.P., J.M., A.P.N, D.J.R., E.D.A.; funding acquisition: E.E.D., D.S.P., M.I., N.S., A.P., A.B., A.S.B., J.D., S.P.; interpreted results: T.V., K.L.B., E.E.D., N.K., M.T., D.S.P., M.I., L.P., J.K., A.T., E.P.; supervision: E.E.D., L.P., D.S.P., N.S., M.I., A.P., A.B., A.S.B.; writing original draft: T.V., K.L.B., E.E.D.; all authors reviewed and edited the manuscript.

## Acknowledgements

We thank members of the Davenport and Parts laboratories for helpful discussion and feedback. In particular, we thank Megan Gozzard, Jacob Hepkema, Matthew Hurles, Seri Kitada, Andrew Lawson, Julie Matte, and Juliane Weller for providing comments on the manuscript and discussing results. We thank the Wellcome Sanger Institute’s Human Genetics Informatics (HGI) team for mapping the bulk RNA-seq reads. This research was funded in whole, or in part, by the Wellcome Trust [Grant numbers 206194 and 220540/Z/20/A]. This study makes use of data generated by the DECIPHER community. A full list of centers who contributed to the generation of the data is available from https://deciphergenomics.org/about/stats and via email from. Funding for the DECIPHER project was provided by Wellcome [grant number WT223718/Z/21/Z]. The Genotype-Tissue Expression (GTEx) Project was supported by the Common Fund of the Office of the Director of the National Institutes of Health, and by NCI, NHGRI, NHLBI, NIDA, NIMH, and NINDS. The data used for the analyses described in this manuscript were obtained from: the GTEx Portal on 09/20/2023. For the purpose of Open Access, the author has applied a CC BY public copyright license to any Author Accepted Manuscript version arising from this submission.

Participants in the INTERVAL randomized controlled trial were recruited with the active collaboration of NHS Blood and Transplant England (https://www.nhsbt.nhs.uk/), which has supported field work and other elements of the trial. DNA extraction and genotyping were co-funded by the National Institute for Health and Care Research (NIHR), the NIHR BioResource (https://bioresource.nihr.ac.uk/) and the NIHR Cambridge Biomedical Research Centre (BRC-1215-20014). RNA-seq was funded as part of an alliance between the University of Cambridge and the AstraZeneca Centre for Genomics Research, and by the NIHR Cambridge Biomedical Research Centre (BRC-1215-20014). The academic coordinating center for INTERVAL was supported by core funding from the NIHR Blood and Transplant Research Unit (BTRU) in Donor Health and Genomics (NIHR BTRU-2014-10024); NIHR BTRU in Donor Health and Behaviour (NIHR203337); UK Medical Research Council (MR/L003120/1); British Heart Foundation (SP/09/002; RG/13/13/30194; RG/18/13/33946); and NIHR Cambridge BRC (BRC-1215-20014; NIHR203312). A complete list of the investigators and contributors to the INTERVAL trial is provided in Di Angelantonio et al.^15^. The academic coordinating center would like to thank blood donor center staff and blood donors for participating in the INTERVAL trial. This work was supported by Health Data Research UK, which is funded by the UK Medical Research Council, Engineering and Physical Sciences Research Council, Economic and Social Research Council, Department of Health and Social Care (England), Chief Scientist Office of the Scottish Government Health and Social Care Directorates, Health and Social Care Research and Development Division (Welsh Government), Public Health Agency (Northern Ireland), British Heart Foundation and Wellcome. The views expressed are those of the authors and not necessarily those of the NIHR or the Department of Health and Social Care.

Personal funding/acknowledgements:

T.V. was supported by a BBSRC iCASE Studentship partly funded by AstraZeneca (BB/V509425/1). A.T. was supported by the Wellcome Trust (PhD studentship 222548/Z/21/Z). E.P. was funded by the EU/EFPIA Innovative Medicines Initiative Joint Undertaking BigData@Heart grant 116074 and is funded by the NIHR BTRU in Donor Health and Behaviour (NIHR203337). J.D. holds a British Heart Foundation Professorship and a NIHR Senior Investigator Award. M.I. is supported by the Munz Chair of Cardiovascular Prediction and Prevention and the NIHR Cambridge Biomedical Research Centre (BRC-1215-20014; NIHR203312). M.I. was also supported by the UK Economic and Social Research Council (ES/T013192/1). A.S.B. has received grants outside of this work from AstraZeneca, Bayer, Biogen, BioMarin and Sanofi.

## Supplementary Information

### Supplementary Figures

**Supplementary Figure 1.** Removal of global expression outliers and inactive gene set validation.

**Supplementary Figure 2.** Misexpression metrics across genes and samples.

**Supplementary Figure 3.** Different properties of misexpressed and non-misexpressed genes across misexpression z-score thresholds.

**Supplementary Figure 4.** Top 75 Human Phenotype Ontology terms underrepresented within misexpressed genes.

**Supplementary Figure 5.** Percentage of misexpression events with a rare SV within 200 kb.

**Supplementary Figure 6.** Enrichment of rare SNV, indels and SVs across genomic windows and misexpression z-score thresholds.

**Supplementary Figure 7.** Enrichment of rare SV classes.

**Supplementary Figure 8.** Enrichment of rare SVs stratified by their class and predicted VEP consequences across misexpression z-score thresholds.

**Supplementary Figure 9.** Properties of misexpression-associated rare SVs.

**Supplementary Figure 10.** Transcriptional readthrough region FPKM and FBNC, transcription readthrough SV consequences and OTP misexpresssion.

**Supplementary Figure 11.** Examples of chimeric misexpression via transcript fusion.

**Supplementary Figure 12.** Intestinal alkaline phosphatase (*ALPI*) misexpression.

### Supplementary tables

**Supplementary Table 1.** *List of 129 inactive genes correlated with any biological or technical covariate*.

**Supplementary Table 2.** *List of 8650 inactive genes*.

**Supplementary Table 3.** *81 gene-level features used for comparing misexpressed and non-misexpressed genes*.

**Supplementary Table 4.** *Genetic variant enrichment results across all 700* enrichments tests.

**Supplementary Table 5.** *23 regulatory features used for comparison of misexpression-associated and control SVs*.

