## Supplementary Figures for "Misexpression of inactive genes in whole blood is associated with nearby rare structural variants"

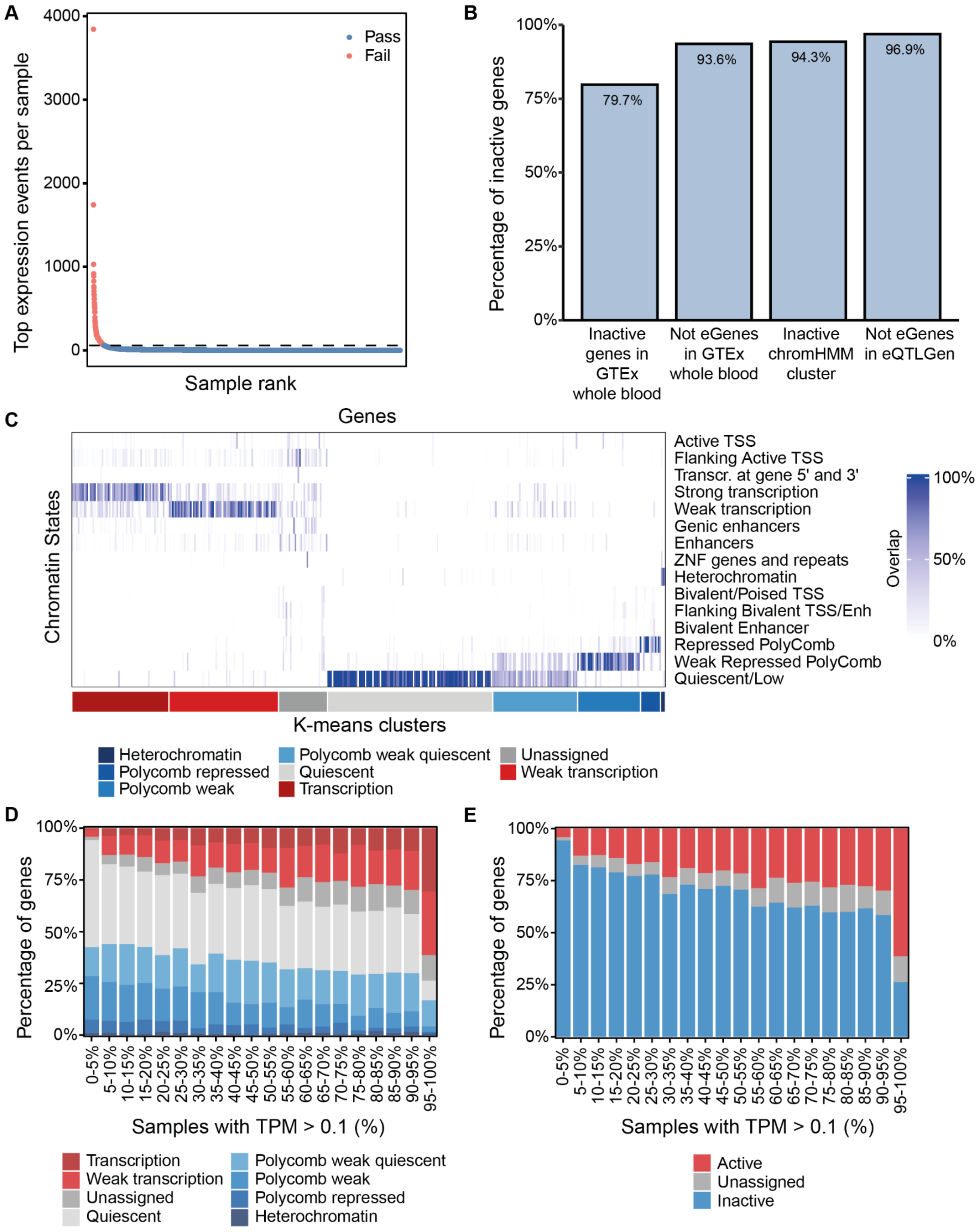


***Supplementary Figure 1. Removal of global expression outliers and inactive gene set validation. A.)*** *Number of top expression events (y-axis) ranked across all samples (x-axis). The dashed line indicates the threshold for removing aberrant samples. Failed samples (red) had a greater number of top expression events than this threshold while samples passing (blue) had a lower number.* ***B.)*** *Different inactive gene validation approaches showing the percentage of inactive genes identified in INTERVAL (y-axis) within different gene sets (x-axis).* ***C.)*** *Heatmap showing the*

*percentage overlap of 60,603 genes (x-axis) over 15 chromHMM states from PBMC data. Genes are clustered into 8 k-means clusters and each cluster is labeled according to the types of overlapping states.* ***D.)*** *Percentage of genes in each k-means cluster stratified by gene expression activity. For each gene, expression activity is quantified as the percentage of samples where the gene has a TPM > 0.1 (x-axis).* ***E.)*** *Percentage of genes labeled as active, inactive or unassigned from chromHMM k-means clusters stratified by gene expression activity. For each gene, activity is quantified as the percentage of samples where the gene has a TPM > 0.1 (x-axis). Inactive genes are defined as having a TPM > 0.1 in less than 5% of samples. GTEx; Genotype-tissue Expression, ZNF; zinc-finger protein. TSS; transcription start site, Enh; enhancer, TPM; transcripts per million.*

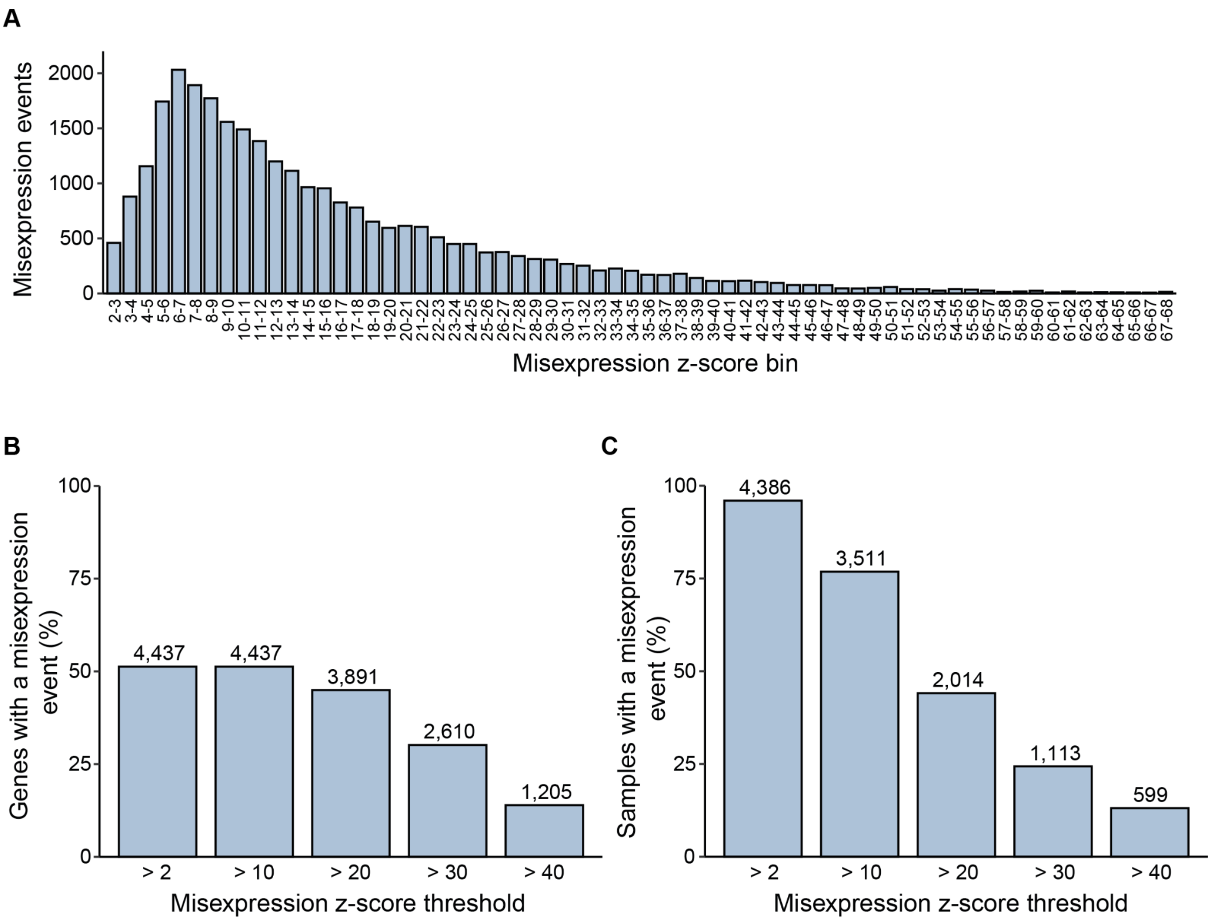


***Supplementary Figure 2. Misexpression metrics across genes and samples.***

***A.)*** *Number of misexpression events across different misexpression z-score threshold bins.* ***B.)*** *Percentage of 8,650 inactive genes that have at least one misexpression event across different misexpression z-score thresholds. Text labels indicate the total number of genes with at least one misexpression event.* ***C.)*** *Percentage of 4,568 samples that have at least one misexpression event across different misexpression z-score thresholds. Text labels indicate the total number of samples with at least one misexpression event.*

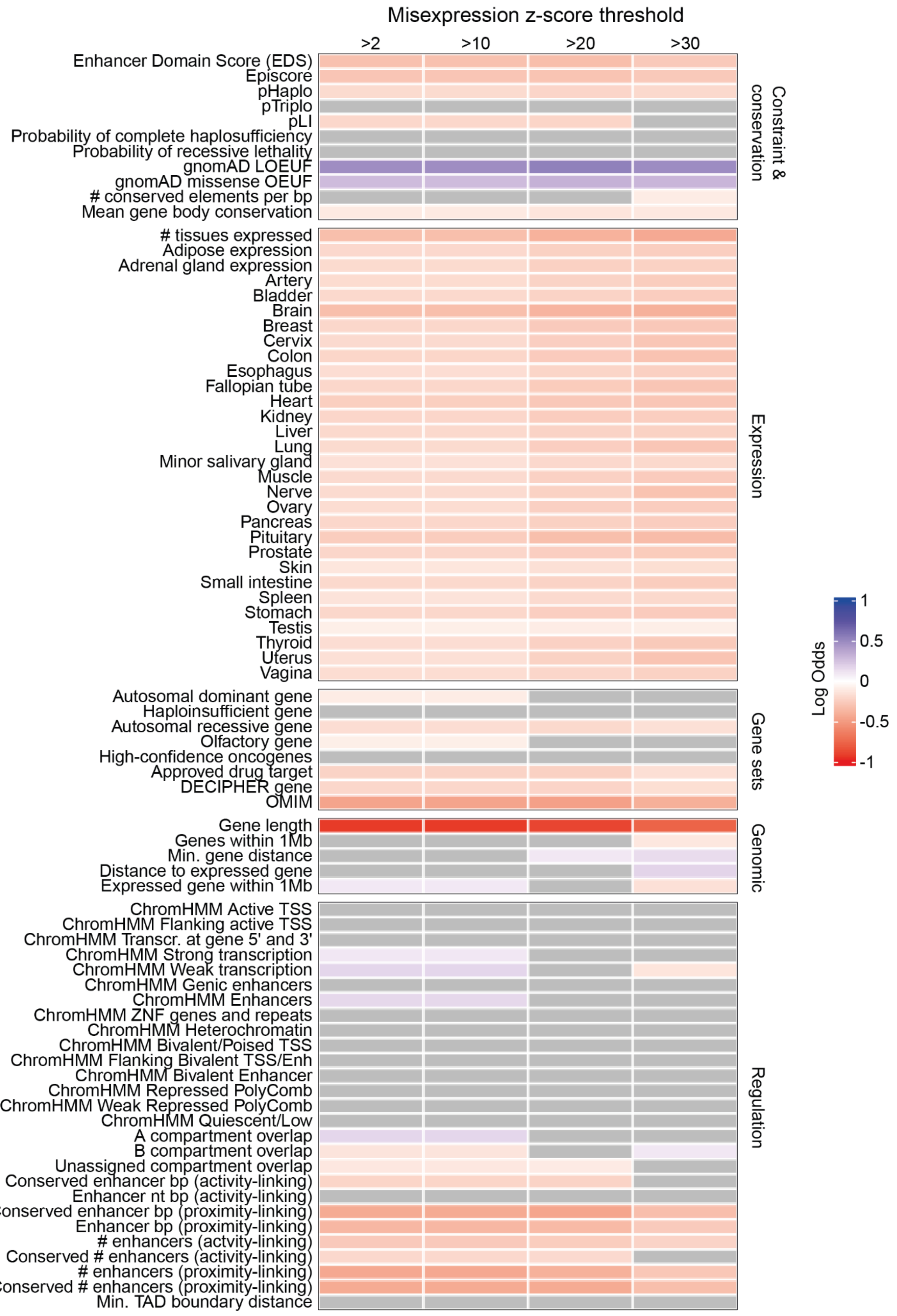


***Supplementary Figure 3. Different properties of misexpressed and non-misexpressed genes across misexpression z-score thresholds.***

*Enrichment of all 81 gene-level features within genes that are misexpressed versus non-misexpressed genes across different misexpression z-score thresholds. Features are grouped* *into different categories. Tiles shaded in gray do not pass a Bonferroni-adjusted p-value threshold ($p < 0.05$). pHaplo; probability of haploinsufficiency, pTriplo; probability of triplosensitivity, pLi; probability of loss-of-function intolerance, LOEUF; loss-of-function observed/expected upper bound fraction, OEUF; observed/expected upper bound fraction, OMIM; Online Mendelian Inheritance in Man, TSS; transcription start site, Enh; enhancer, TAD; topologically associating domain, ZNF; zinc-finger protein.*

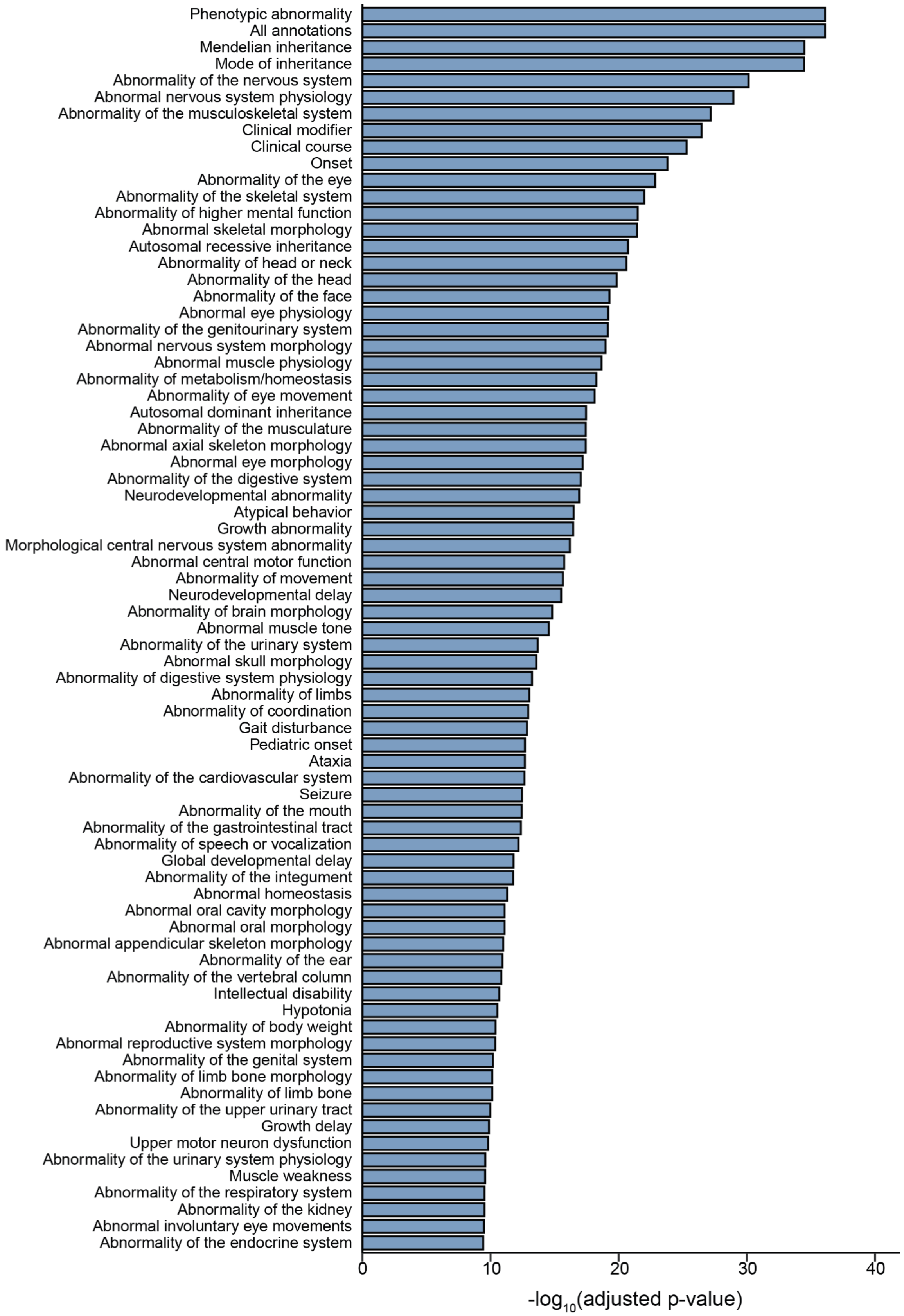


***Supplementary Figure 4. Top 75 Human Phenotype Ontology terms underrepresented within misexpressed genes.***

*Top 75 HPO terms, by -log$_{10}$(adjusted p-value) on the x-axis, underrepresented within 4,437 misexpressed genes using all 8,650 inactive genes as the custom background.*

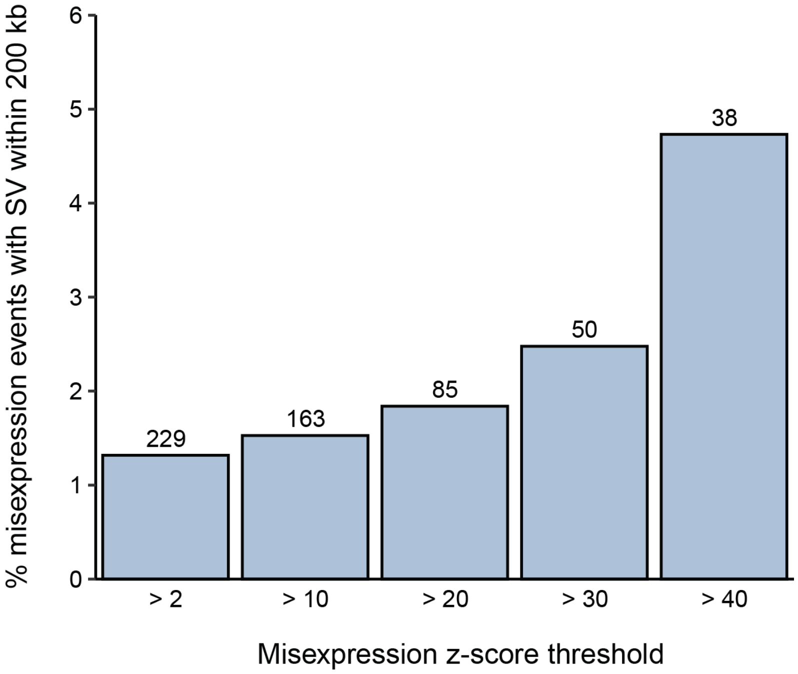


***Supplementary Figure 5. Percentage of misexpression events with a rare SV within 200 kb.*** *Percentage of misexpression events (y-axis) with a rare SV within 200 kb at different misexpression z-score thresholds (x-axis). Text labels indicate the total number of misexpression events with an SV.*

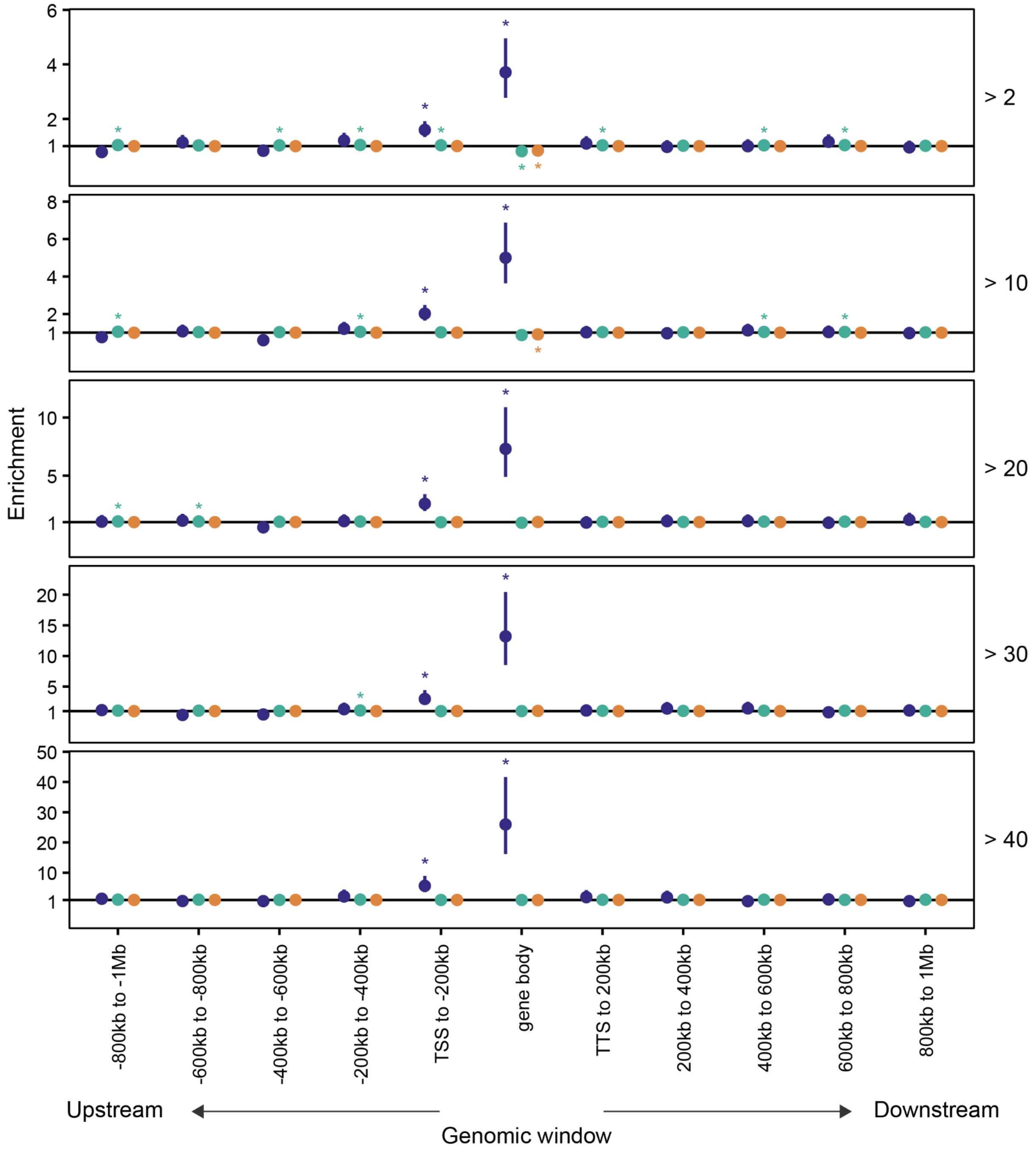


***Supplementary Figure 6. Enrichment of rare SNV, indels and SVs across genomic windows and misexpression z-score thresholds.***

*Enrichment of rare (MAF < 1%) SNVs (orange), indels (green) and SVs (blue) within 200 kb genomic windows and the body of the misexpressed gene across different misexpression z-score thresholds. Enrichments were calculated as the relative risk of having a nearby rare variant type given the misexpression status. The line at enrichment = 1 indicates no enrichment; Asterisks positioned either side of the line indicate significant enrichment or underenrichment after Bonferroni correction. Bars represent 95% Wald confidence intervals of the relative risk estimates. SNV; single nucleotide variant, SV; structural variant, TTS; transcription termination site, TSS; transcription start site.*

A

B

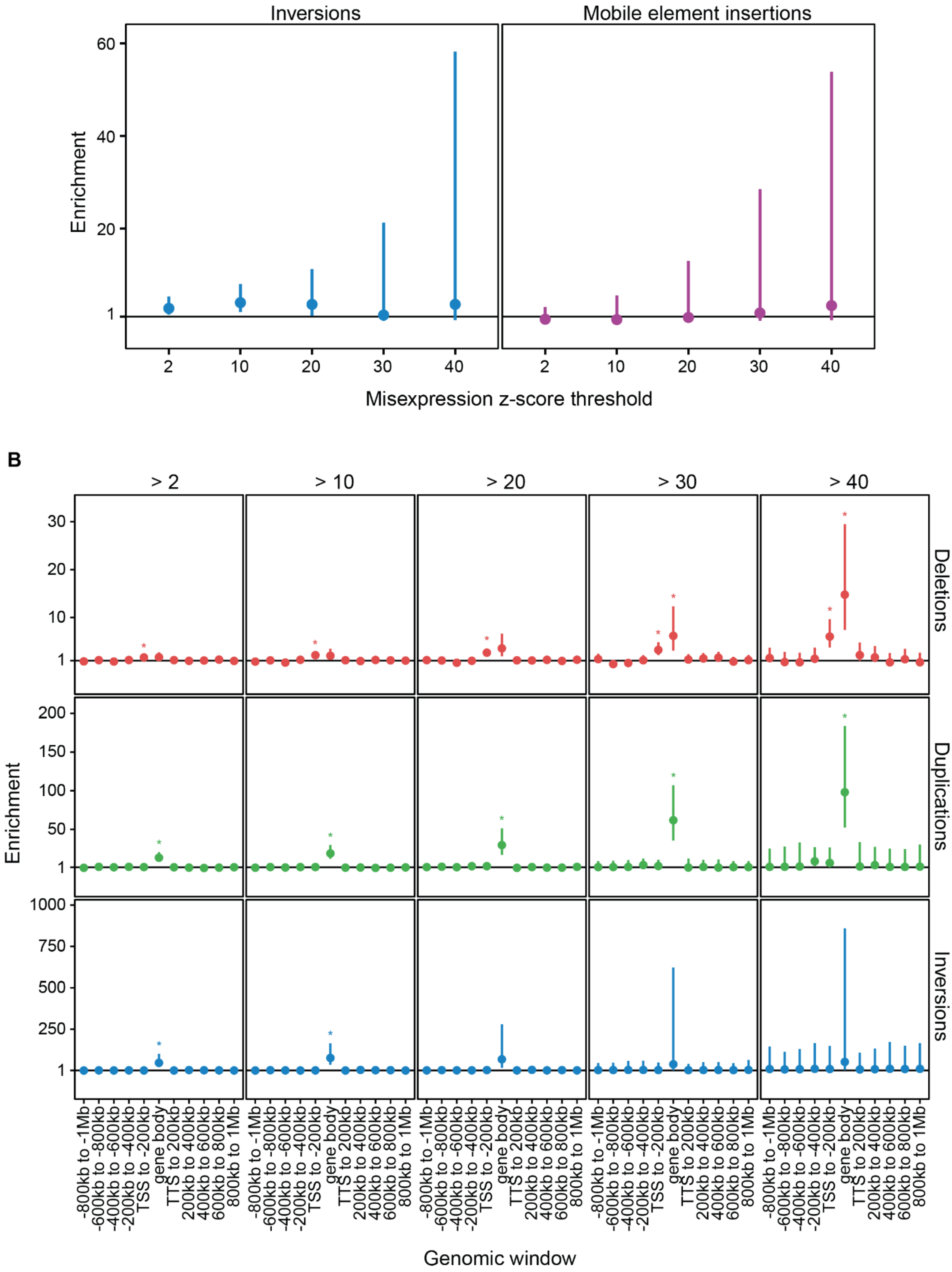


***Supplementary Figure 7. Enrichment of rare SV classes.***
*Enrichments were calculated as the relative risk of having a nearby variant type or consequence given the misexpression status. Bars represent 95% Wald confidence intervals of the relative risk estimates. The line at enrichment = 1 indicates no enrichment; stars positioned either side of the line indicate significant enrichment or underenrichment after Bonferroni correction.* ***A.)*** *Enrichment of rare (MAF < 1%) inversions and mobile element insertions in a ±200 kb window around the tested genes across*

*different misexpression z-score thresholds.* ***B.)*** *Enrichment of rare (MAF < 1%) deletions, duplications, and inversions within 200 kb genomic windows and the body of the misexpressed gene across different misexpression z-score thresholds. TSS; transcription start site, TTS; transcription termination site.*

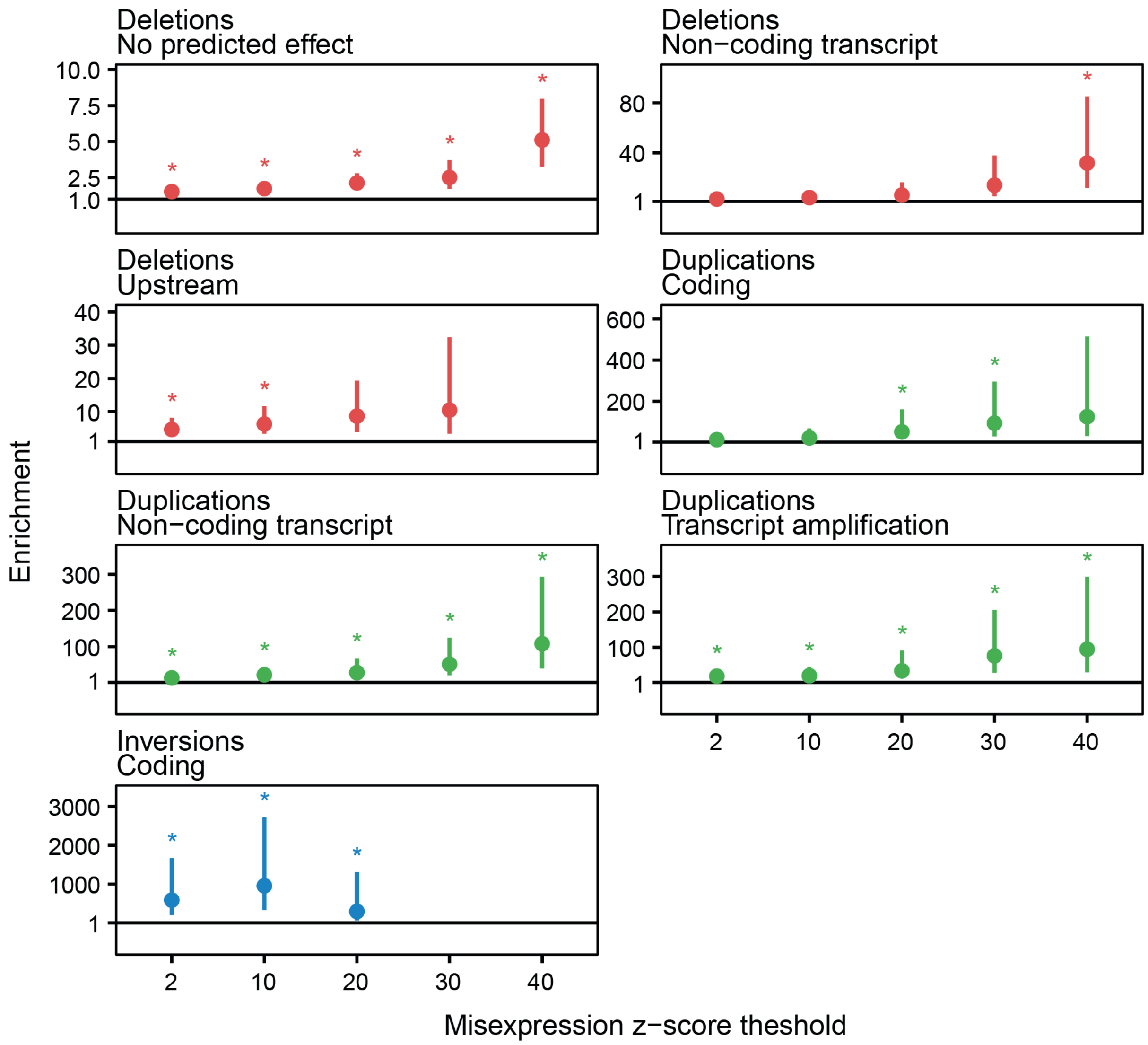


***Supplementary Figure 8. Enrichment of rare SVs stratified by their class and predicted VEP consequences across misexpression z-score thresholds.***

*Enrichments were calculated as the relative risk of having a nearby variant consequence given the misexpression status. The line at enrichment = 1 indicates no enrichment; asterisks positioned either side of the line indicate significant enrichment or underenrichment after Bonferroni correction. Bars represent 95% Wald confidence intervals of the relative risk estimates. Only SV consequences with at least one Bonferroni significant enrichment at any z-score threshold are shown. Missing points indicate tests failing to pass the nominal p-value threshold (p ≥ 0.05).*

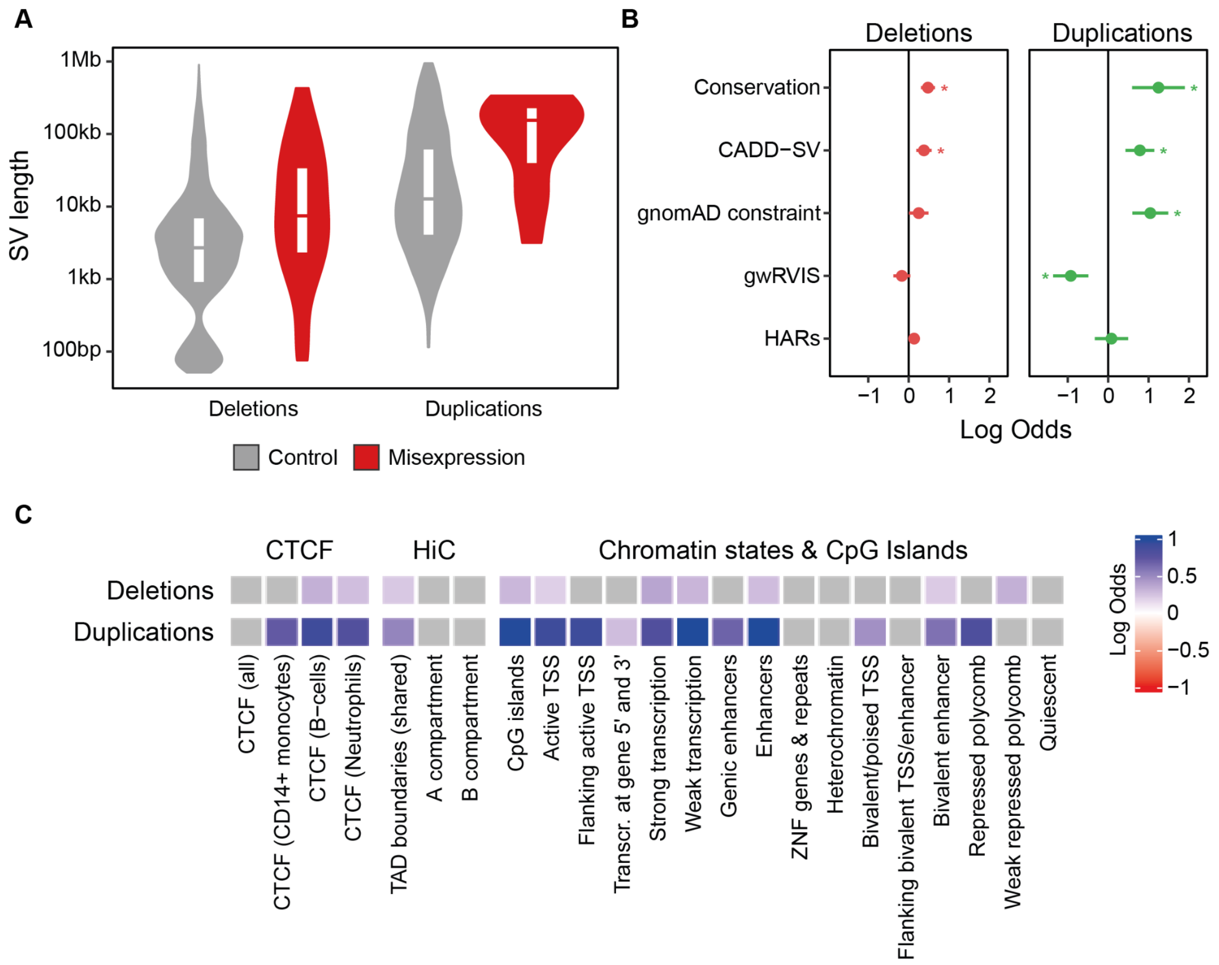


***Supplementary Figure 9. Properties of misexpression-associated rare SVs.***
***A.)*** *SV length distributions of misexpression-associated and control duplications and deletions restricted to singletons only.* ***B.)*** *Enrichment (x-axis) without adjusting for SV length of misexpression-associated deletions (left panel, red) and duplications (right panel, green) compared to controls for genomic scores (y-axis) including evolutionary conservation (phyloP), predicted deleteriousness (CADD-SV), constraint (gnomAD z-score constraint and gwRVIS), and HARs. Enrichments were calculated as the log odds ratio with lines indicating 95% confidence intervals for the fitted parameters using the standard normal distribution. Asterisks indicate significant enrichment after Bonferroni correction.* ***C.)*** *Enrichment without adjusting for SV length of misexpression-associated deletions and duplications compared to controls for regulatory features including CTCF candidate cis-regulatory elements from ENCODE, TAD boundaries shared across multiple cell-lines, A and B compartments, chromatin states from the Roadmap Epigenomics Project and CpG islands from the UCSC genome browser. Enrichments were calculated as the log odds ratio and tiles shaded in gray do not pass Bonferroni correction. CADD-SV; combined annotation dependent depletion for SVs, gwRVIS; genome-wide residual variation intolerance score, TADs; topologically-associated domains, HARs; human accelerated regions, TSS; transcription start site, CTCF; CCCTC-binding factor, ZNFs; zinc-finger proteins.*

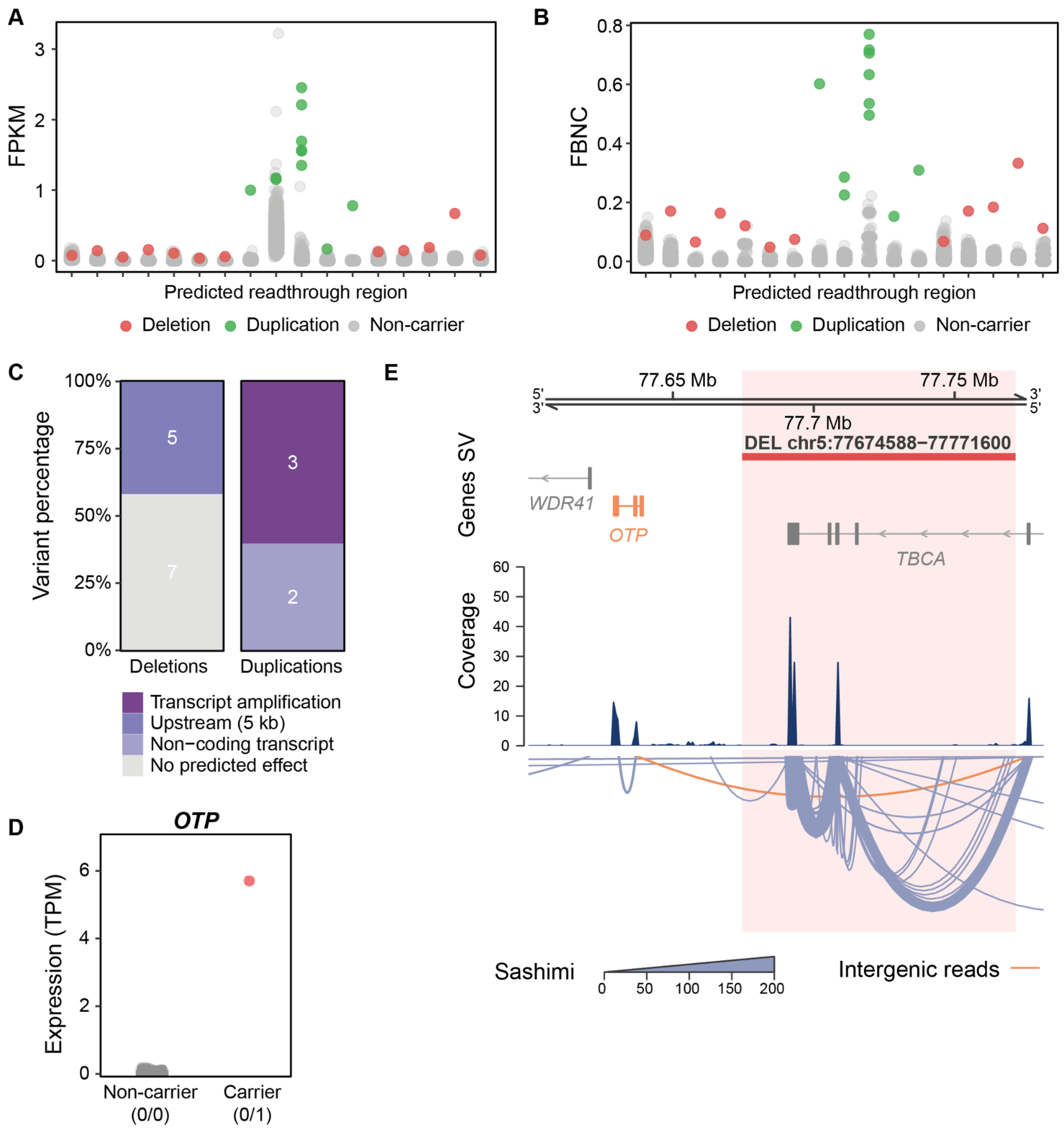


***Supplementary Figure 10. Transcriptional readthrough region FPKM and FBNC, transcription readthrough SV consequences and OTP misexpresssion.***

***A.)*** *FPKM and* ***B.)*** *FBNC at the 17 predicted readthrough regions for carriers of candidate transcriptional readthrough deletions (red) and duplications (green), as well as non-carriers (gray).* ***C.)*** *Proportion of candidate transcriptional readthrough deletions and duplications by their predicted VEP consequence on the misexpressed gene.* ***D.)*** *Expression of OTP in a DEL chr5:77674588−77771600 carrier and non-carriers. Red color indicates samples passing the misexpression threshold TPM > 0.5 and z-score > 2 while gray samples are below this threshold.* ***E.)*** *Deletion of the 3' end of TBCA results in transcriptional readthrough. Transcriptional readthrough leads to OTP misexpression (orange gene) and intergenic splicing between TBCA and OTP (intergenic reads, orange). In the sashimi plot, the line width corresponds to the number of reads spanning a given junction. FPKM; fragments per kilobase of transcript per million mapped reads, FBNC; fraction of bases with non-zero coverage. TPM; transcripts per million.*

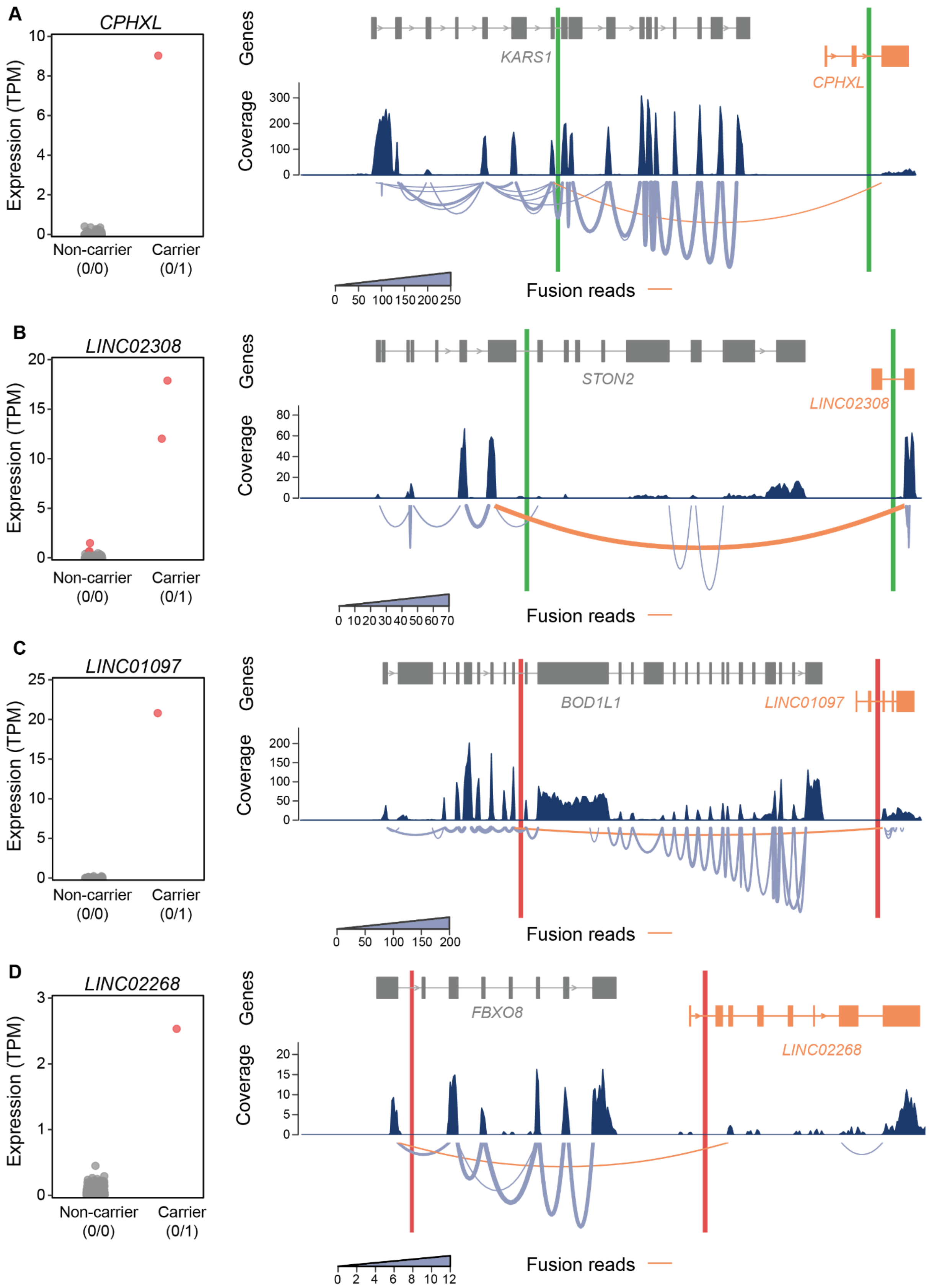


***Supplementary Figure 11. Examples of chimeric misexpression via transcript fusion.***

*Comparison of carrier and non-carrier expression with FusionInspector visualization of the fusion transcript in a carrier for **A.)** CPHXL misexpression via KARS–CPHXL fusion in DUP chr16:75636427-75717471 carrier, **B.)** LINC02308 misexpression via STON2–LINC02308 fusion in*

*DUP chr14:81376672-81444942 carrier,* ***C.)*** *LINC01097 misexpression via BOD1L1–LINC01097 fusion in DEL chr4:13529219-13608506 carrier, and* ***D.)*** *LINC02268 misexpression via FBXO8–LINC02268 fusion in DEL chr4:174159287-174273823 carrier. Red color indicates samples passing the misexpression threshold TPM > 0.5 and z-score > 2 while gray samples are below this threshold. In the sashimi plot, the line width corresponds to the number of reads spanning a given junction. The misexpressed gene and fusion reads are colored in orange. Deletion and duplication breakpoints are colored in red and green, respectively. Introns have been shortened for visualization and breakpoint positions have been approximated accordingly. TPM; transcripts per million.*

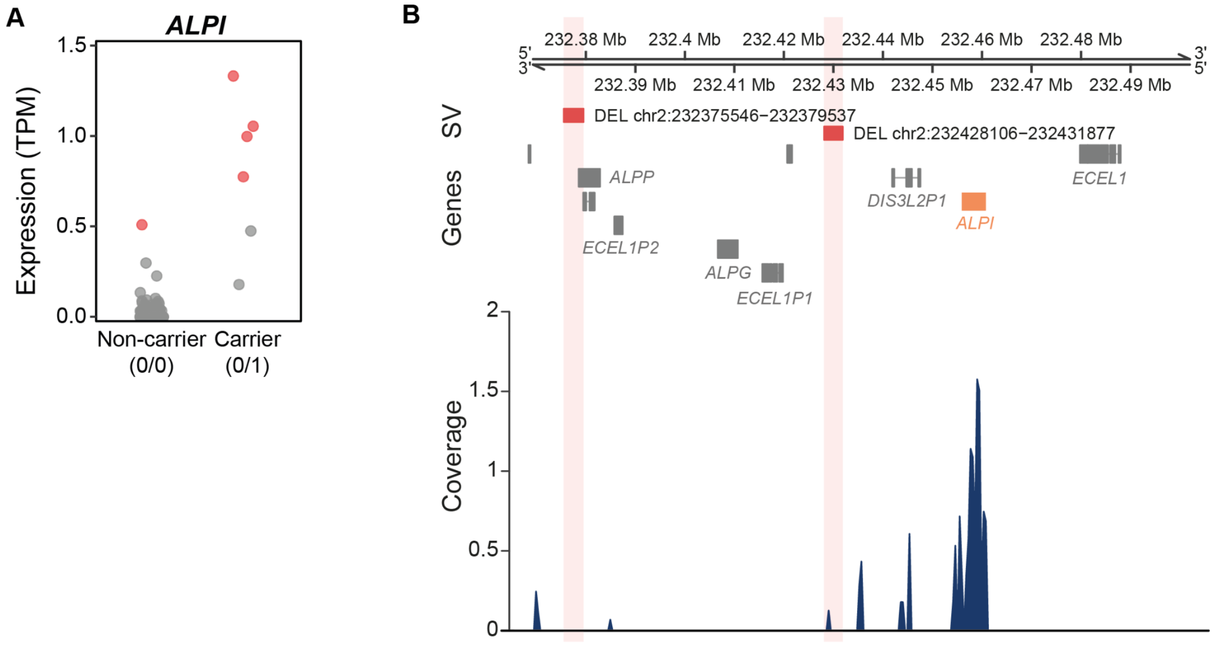


***Supplementary Figure 12. Intestinal alkaline phosphatase (ALPI) misexpression.***
***A.)*** *Expression of ALPI in DEL chr2:232375546-232379537 and DEL chr2:232428106-232431877 carriers and non-carriers. Red color indicates samples passing the misexpression threshold TPM > 0.5 and z-score > 2 while gray samples are below this threshold.* ***B.)*** *Position of DEL chr2:232375546-232379537 and DEL chr2:232428106-232431877 relative to the misexpressed gene ALPI (orange gene). Deletions are marked in red. TPM; transcripts per million.*
